# CRISPR-engineered inducible flocculation in *Komagataella phaffii* enables enhanced biomass separation for biopharmaceutical production

**DOI:** 10.64898/2026.02.05.704028

**Authors:** Elena Ivanova, Paul Ramp, Natalie Zimmer, Markus Mund, Elena Antonov, Christoph Schiklenk, Daniel Degreif

## Abstract

Biomass separation represents a critical bottleneck in *Komagataella phaffii*-based biopharmaceutical processes, as typically high cell densities of 40 – 50 % create significant operational, technical and economic challenges for harvest operations. Yeast cell aggregation (flocculation) provides a solution to accelerate cell sedimentation by increasing particle size, thus allowing to improve biomass-supernatant separation efficiency during both natural gravity settling and (continuous) centrifugation operations. This study demonstrates successful engineering of *K. phaffii* strains with an inducible flocculation phenotype using CRISPR/Cas9-based genome editing to integrate the *Saccharomyces cerevisiae FLO1* (*ScFLO1*) gene under control of various regulatory elements, including methanol-inducible and derepressible promoters. Flocculation strength could be enhanced by implementing transcriptional positive feedback circuits based on the methanol-inducible *AOX1* promoter. To address methanol-free production requirements, we developed alternative systems to retrofit P*_AOX1_*-based *ScFLO1* expression and exploited the derepressible PDF promoter, offering broader compatibility with biopharmaceutical manufacturing facilities. Flocculating cells cultivated in a bioreactor demonstrated significantly improved sedimentation behavior, with considerably lower supernatant turbidity after short low-speed centrifugation compared to non-flocculating controls. Crucially, cell flocculation had no negative impact on product amount and quality when expressing a multivalent NANOBODY^®^ VHH molecule with pharmaceutical relevance. Thus, this work establishes the first genetically engineered flocculation system in *K. phaffii* compatible with recombinant protein production, providing the basis for an innovative approach to streamline harvest operations in biopharmaceutical processes.

## 1. Introduction

*Komagataella phaffii* (formerly known as *Pichia pastoris*) has emerged as a versatile and powerful eukaryotic expression system for biopharmaceutical production. Its ability to grow to high cell densities, coupled with the availability of strong and tightly regulated promoters such as the methanol (MeOH)-inducible *AOX1* promoter (Vogl and Glieder, 2013) or recently established MeOH-free expression systems (Groeve et al., 2023; Shirvani et al., 2019; Vogl et al., 2018), enables high-level expression of heterologous genes. With that, this methylotrophic yeast combines several advantages of prokaryotic and higher eukaryotic hosts, making it particularly suitable to produce complex recombinant proteins.

The inherent advantages of *K. phaffii* as an expression host are rooted in its remarkable capabilities for protein processing. These capabilities have even already been subject to extensive optimization through several strain engineering endeavors. Notably, significant advancements have been achieved in the realm of glycosylation, where modifications have been implemented to emulate humanized glycosylation patterns (Beck et al., 2010; Laukens et al., 2015). Furthermore, substantial progress has been made in enhancing disulfide bond formation and augmenting protein folding efficiency through the co-expression of auxiliary proteins and the activation of molecular chaperones (Gasser et al., 2006; Groeve et al., 2023; Guerfal et al., 2010; Navone et al., 2021). Such refined post-translational processes are crucial as they are intrinsically linked to the biological activity and structural stability of a wide array of therapeutic proteins. Another particularly important feature of *K. phaffii* is its large capacity to secrete recombinant proteins directly into the culture medium, greatly simplifying purification. The use of well-characterized signal peptides, such as the alpha-mating factor prepro-leader from *Saccharomyces cerevisiae* and derivatives, enable efficient targeting of proteins to the secretory pathway (Lin-Cereghino et al., 2013; Zou et al., 2022).

In production processes based on secretory protein expression, harvest operations need to efficiently separate yeast biomass from the product-containing culture supernatant. At industrial scale, such harvest operations typically involve continuous centrifugation-based separation methods (e. g. disc-stack centrifugation) to remove biomass followed by multi-stage filtration (as a combination of depth- and sterile-filtration). These steps are required to generate product containing, particle-free solution that is suitable for chromatography purification steps (Depth Filtration vs. Centrifugation, 2014; Wang et al., 2006). In high performing bioprocesses, *K. phaffii* high-cell-density fermentations can reach up to 40% - 50% (w/v) total biomass (Cregg et al., 2000; Liu et al., 2020; Shemesh and Fishman, 2024), hence the separation of the biomass by such harvesting regimes poses significant technical and economic challenges. Single-pass disc-stack centrifugation may not achieve the desired separation at high cell densities, necessitating multiple passes or dilution of the cell broth. Both options introduce disadvantages for the process, such as significantly increased process intermediate volumes or significantly extended process times, which bring challenges with regard to process intermediate hold times. Furthermore, continuous centrifugation processes are considered technically complex, and the lack of suitable small-scale models limit process development options. Therefore, the clarification steps at industrial scale often lack efficiency and robustness. To compensate for such effects and to efficiently deplete host cell proteins or debris released by co-secretion or through cell lysis, multi-stage depth filtration with expensive filter materials is required, which significantly contributes to overall process costs (Felo et al., 2013). While these technical setups can deliver suitable clarification levels for downstream processes, they typically only achieve harvest yields of approximately 80 %, thus limiting overall process yields. Only intensive, process-specific harvest optimization achieves yields of up to 90 % (Hebbi et al., 2025).

Given such limitations, the biopharmaceutical industry is exploring alternatives to traditional clarification strategies by making use of cell precipitation and flocculation. These techniques are routinely applied at large scale in other industries such as brewing (Gassara et al., 2015) or wastewater treatment (Lee et al., 2014). Among these approaches, processes using acid-or salt-based precipitation methods are favorable options due to their compatibility with downstream process steps. In mammalian processes for monoclonal antibody production, cationic polymers such as chitosan and polydiallyldimethylammonium chloride (PDADMAC) have already been successfully implemented as chemical flocculants to remove host cell DNA and proteins (McNerney et al., 2015). Also, for *K. phaffii*-based processes, flocculation-based clarification by addition of inexpensive and biologically compatible salts has been described (Hebbi et al., 2025). However, addition of chemical flocculants introduces additional raw materials into processes which need to comply with high standards for biopharmaceutical production and regulatory guidelines require the demonstration of quantitative flocculant removal.

A promising, alternative approach to facilitate cell-liquid separation is yeast flocculation - a naturally occurring, reversible, non-sexual aggregation of yeast cells into multicellular clumps or flocs. According to Stokes’ law, the sedimentation velocity of a spherical particle is proportional to the square of its radius (r²). Consequently, cell flocs with increased radii exhibit accelerated sedimentation compared to individual cells, thus enhancing biomass-supernatant separation efficiency under identical gravitational field conditions (Soares, 2009). This natural trait, extensively studied in *S. cerevisiae* and related species, offers a potential for a low-cost, scalable, and sustainable alternative to chemical flocculation and could enhance efficiency of mechanical separation methods. Classical yeast flocculation is mediated by cell-wall anchored lectin-like proteins, called flocculins (encoded by *FLO* genes), which bind cell wall mannose residues of adjacent cells, thus mediating cell-cell aggregation (Soares, 2011) (Fig. 1a). The *FLO* gene repertoire varies between different *S. cerevisiae* strains and genes are activated by different factors and triggers. *FLO* genes are induced by nitrogen starvation or adverse environmental conditions like elevated ethanol concentrations, and temperature stress (Verstrepen et al., 2003). These typically act via an Flo8p-dependent induction pathway (Kobayashi et al., 1996). In addition, Flo8-independent induction of Flo1p-mediated flocculation upon hypoxic conditions has been recently described (Degreif et al., 2017).

**Fig. 1:**
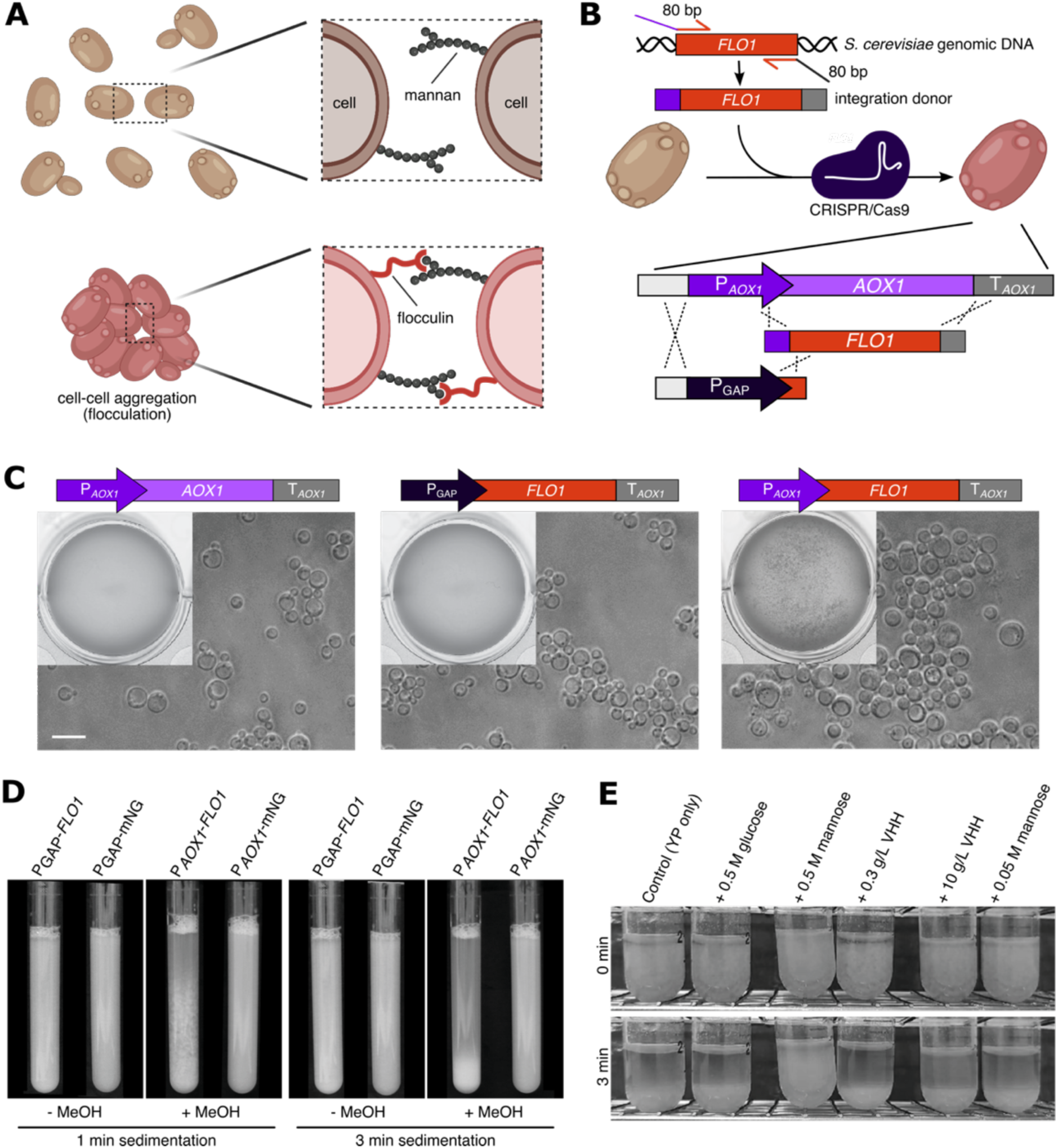
Engineering *Sc*Flo1p-mediated flocculation. **A:** Molecular mechanism of flocculation: Flocculins exposed on the cell surface bind mannans present in the cell walls of other cells, thus mediating cell-cell aggregation. Created in BioRender. Degreif, D. (2026) https://BioRender.com/54nbqy4 **B:** Schematic representation of CRISPR/Cas9-mediated engineering of flocculating *K. phaffii* strains. The *FLO1* gene was amplified from *Saccharomyces cerevisiae* genomic DNA by PCR and used as integration donor. The *ScFLO1* donor was in a first step integrated into the *AOX1* locus generating the P*_AOX1_*-*ScFLO1* construct. In a second step, the *AOX1* promoter (P*_AOX1_*) was replaced by the GAP promoter (P_GAP_). Created in BioRender. Degreif, D. (2026) https://BioRender.com/54nbqy^4^ **C:** Microscopic and macroscopic comparison of the floc size of indicated strains. For macroscopic analysis, cell suspension was transferred into a well of a 24-well plate (top left corner). Scale bar = 10 µm. **D:** Sedimentation of indicated strains grown in shake flask under specified conditions (-MeOH, +MeOH) after transfer into culture tubes. Sedimentation was photographically documented after 1 min and 3 min. **E:** Competitive flocculation inhibition experiments. Indicated substances (D-glucose, D-mannose or cultivation medium containing 10 g/L NANOBODY VHH A) were added to a suspension of the flocculating P*_AOX1_*-*ScFLO1* (yEIA010) cells.

In contrast, induction of endogenous *FLO* gene *in K. phaffii* occurs under conditions of slow growth rates and/or carbon source limitation (De et al., 2020) and promote pseudohyphal growth. This pseudohyphal growth is similar to filamentous growth induced by *FLO11* expression in *S. cerevisiae* (Lambrechts et al., 1996) rather than a cell-cell-adhesion/flocculation phenotype supporting sedimentation, which is mediated in *S. cerevisiae* by the *FLO1*, *FLO5*, *FLO9*, *FLO10* family (Govender et al., 2008; Teunissen and Steensma, 1995).

In this study, we explore cell adhesion-based flocculation of *K. phaffii* for bioprocesses and assess its potential to facilitate biomass separation from product containing culture supernatant to streamline harvest operations. Given the absence of native *FLO* genes in *K. phaffii* mediating classical cell-cell-adhesion, we engineered strains capable of flocculation through the heterologous expression of the flocculation-associated gene *FLO1* derived from *S. cerevisiae*.

Our experimental approach was informed by two key studies demonstrating successful expression of *ScFLO1* in *K. phaffii*. The first study achieved high-level expression of the ScFlo1p *N*-terminal domain for protein characterization (Goossens et al., 2011). More recent work demonstrated expression of functional *ScFLO1* in *K. phaffii* KM71 using the constitutive glyceraldehyde-3-phosphate dehydrogenase promoter (P_GAP_). In that study, the *ScFLO1* gene, amplified from *S. cerevisiae* S288c genomic DNA, was cloned into a *K. phaffii* expression vector and randomly integrated into the genome. Successful integration was verified both through PCR analysis and by the observation of a flocculation phenotype, which provided protection against stressors such as D-lactic acid accumulation (Sae-Tang et al., 2023).

However, to our knowledge, *K. phaffii* strains have not been engineered for inducible flocculation in a manner compatible with the production of biopharmaceutical-relevant heterologous proteins.

Accordingly, this study sought to establish a strain engineering framework to equip *K. phaffii* with a flocculation phenotype applicable in an industrial recombinant protein production setting. Emphasis was placed on both strain construction and bioreactor cultivation. Using these engineered strains, we evaluated the feasibility of employing genetically induced flocculation as a cell clarification strategy, quantified performance, and assessed the compatibility with conventional mechanical clarification methods.

## 2. Materials and methods

### 2.1. Plasmid construction

For the generation of plasmid pBSY5Z_NANOBODY^®^ VHH B, the plasmid pBSY5Z (bisy GmbH, Graz), which confers resistance to the antibiotic Zeocin, was used as starting construct. The NANOBODY^®^ VHH B coding sequence was synthetized by GeneArt (Thermo Fisher Scientific) as a synthetic DNA construct downstream of and in frame with the alpha mating factor (aMF) signal peptide sequence from *S. cerevisiae* and was delivered cloned into a pMA-RQ plasmid backbone. Both plasmids were digested with SapI (NEB) according to manufacturer’s instructions to remove the stuffer sequence (pBSY5Z) or to release the DNA fragment encoding the aMF-NANOBODY^®^ VHH B fusion protein. pBSY5Z plasmid backbone (3068 bp) and aMF-NANOBODY^®^ VHH B encoding DNA fragment (2148 bp) fragments were purified by gel extraction (QIAquick gel extraction kit, Qiagen) and assembled by ligation (Quick ligation kit, NEB).

Plasmids pBB3Zi_086, pBB3Hi_095, pBB3Hi_095, and pBB3Zi_098 were generated via Golden Gate assembly as described (Prielhofer et al., 2017). pBB1 basic Golden Gate modules (pBB1_PDF_12, pBB1_PDC_12, pBB1_MIT1_23, pBB1_MXR1_23, pBB1_ScRPL3TT_34) were synthetized from GeneArt (Thermo Fisher Scientific). The entry vector backbones (pBB3) mediating Zeocin-(Zi) and Hygromicin (Hi)-resistance were cloned from pBSYBiE1 (Zi) and pBSYBiE1H (Hi) (bisy GmbH). These were SmiI/BamHI-digested and the acceptor cassette (synthetized by GeneArt, Thermo Fisher Scientific) was inserted by Gibson Assembly (NEBuilder, NEB) to generate pBB3Zi_14 and pBB3Hi_14.

Plasmids pBB1_mNG_23 and pBB1_PGAP_12 that served as templates for donor DNA generation by PCR were ordered from GeneArt (Thermo Fisher Scientific).

Cloning of the CRISPR-Cas9 plasmids is described below in the separate section. All oligonucleotides used for cloning and generation of donor DNA are listed in Tab. S1.

All plasmids used and generated in this study are listed in Tab. S2.

### 2.2. Yeast strains and strain construction

All yeast strains in this study are derivates of NRRL Y-11430 or NRRL Y-11430 + *HAC1*, a *Komagataella phaffii* NRRL Y-11430 strain overexpressing the endogenous Hac1 protein under control of the derepressible *CTA1*/*CAT1* promoter (Groeve et al., 2023).

For preparation of electrocompetent *K. phaffii* cells, 20 mL of YPD medium in a 100 mL shake flask were inoculated with the respective strain and cultured overnight at 28 °C and 180 rpm. From this preculture, 50 mL YPD medium in 300 mL shake flask was inoculated to OD_600_ of ∼ 0.2. The main culture was grown at 28 °C for 4.25 h until an OD_600_ of ∼ 0.8 was reached. Cells were pelleted by centrifugation at 500 x g for 5 min at 4 °C. The supernatant was discarded, and the cells were resuspended in 9 mL ice-cold BEDS-solution (1 M sorbitol, 10 mM bicine; 3% ethylene glycol, 5% DMSO) supplemented with 1 mL 1 M DTT. This suspension was incubated for 5 min at room temperature (RT). Subsequently, 800 µL aliquots were prepared and frozen at -80 °C until use for transformation.

For electroporation, an aliquot of competent cells was thawed on ice. Yeast cells were sedimented by centrifugation (500 x g, 5 min, 4 °C). The supernatant was aspirated by pipetting and discarded. Cells were resuspended in 80 µL ice-cold BEDS-solution and the cell suspension was transferred to an electroporation cuvette and mixed with desalted DNA. The cell-DNA mixture was incubated on ice for a 2 - 5 min. Electroporation was performed at 2 kV, 25 µF and 200 Ω. Immediately following the electronic pulse, a 1:1 mixture of 1 M sorbitol and YPD medium was added. Cells were incubated at 28 °C for 2 h shaking at 120 rpm to recover in a 50 mL conical tube. Aliquots of the culture were plated out on YPD agar plates supplemented with hygromycin (200 µg/mL) for selection and incubated at 28 °C for 3 days.

All yeast strains used and generated in this study are listed in Tab. S3.

### 2.3. CRISPR-Cas9 assisted *K. phaffii* genome manipulations

Genome engineering was based on a modified version of the previously published CRISPR-Cas9 plasmid R_C2_gRNAgut1_kan (Dalvie et al., 2020), in which the anti-*GUT1* sgRNA was exchanged by a SapI-site-flanked *E. coli ccdB* counterselection marker. This design allows for oligo-based cloning of sgRNA target sequences (Mund et al., 2023) (Fig. S1a). In addition, the antibiotic resistance was changed to hygR. The sequence encoding the SapI-flanked *ccdB* stuffer was amplified by PCR (Q5, NEB) using primers O56 & O57. The R_C2_gRNAgut1_kan backbone was PCR-amplified as two fragments using primer pairs O52 & O53 and O54 & O55. The three PCR products were assembled by Gibson Assembly (NEBuilder, NEB), resulting in pCas9Gc_ccdB.

To exchange the selection marker in pCas9Gc_ccdb, the plasmid backbone was amplified by PCR (Q5, NEB) as two fragments using primer pairs O53 & O58 and O55 & O59. The sequence encoding hygromycin resistance marker was ordered from GeneArt (Thermo Fisher). The stuffer sequence was amplified by PCR (Q5, NEB) using primers O60 & O61. The three PCR products were assembled by Gibson Assembly (NEBuilder, NEB) according to manufacturer’s instructions. The resulting plasmid was named pCas9Hc_ccdB.

To introduce sgRNA target sequences into pCas9Hc_ccdB, forward and reverse oligonucleotides encoding the target sequence (10 pmol each) and respective overhangs were hybridized in 50 μL annealing buffer (50 mM NaCl, 10 mM Tris, 1 mM EDTA) by heating to 95 °C for 5 min followed by 0.1 °C/s cooling to 4 °C in a thermocycler. 1 μL of ds-oligo solution was ligated into 50 ng SapI-digested pCas9Hc_ccdB using T4 DNA Quick Ligase (NEB). Following oligonucleotides (see Tab. S1) were respectively annealed to address the indicated loci: O39 & O40 (*AOX1*), O41 & O42 (P*_AOX1_*), O43 & O44 (int6 locus), O45 & O46 (int18 locus), O47 & O48 (PNSII-4 locus), O49 & O50 (PNSI-2 locus).

Donor DNA for site-specific genomic integration was generated by PCR from template DNA with primer overhangs adding 80 bp homology sequences to the PCR product for homology-directed integration. Tab. S4 lists the primers and template DNAs. PCR products were desalted by silica column purification (QIAquick, QIAGEN).

To introduce genomic modifications, electrocompetent *K. phaffii* strains were transformed by electroporation with donor DNA (> 1 µg) and 300 ng of the pCas9Hc plasmid targeting the respective locus. Tab. S5 lists starting strains and CRISPR system components used for strain engineering.

Genomic integrations and deletions were verified by colony PCR using primers (Tab. S5) that amplify the borders resulting from the recombination event. Colony material was resuspended in 20 µL water and heated to 95 °C for 10 min in a thermocycler. 2 µL of the suspension were used as template. Elongation times were adjusted to the expected amplicon length in case of successful genome modification. Genomic DNA was extracted from positive clones (Puregene Cell Kit, QIAGEN) and used as template for a PCR using primers (Tab. S5) that amplify the full-length integrated DNA element beyond all recombination sites. The PCR product was purified (QIAquick, QIAGEN) and subjected to Oxford Nanopore-based sequence verification (PlasmidEZ, Azenta).

### 2.4. Cultivation in 24-well deep well plates

For small-scale expression tests, strains were cultured in 24-well deep well plates (Riplate, Ritter Plastic) in 2 mL complex media (10 g/L yeast extract, 10 g/L peptone, 1 % (w/v) glycerol) at 28 °C for 72 h (80 % humidity, 280 rpm, Kuhner shaker). For constructs containing MeOH-inducible promoters, flocculation or mNeonGreen (mNG) expression was induced by the addition of 1 % (v/v, final concentration) MeOH after 72 h. Samples for fluorescence measurements and Bradford assay were collected after 72 h (day 3) and 96 h (day 4). Samples for visual floc size comparison were collected after 75 h, 78 h or 96 h.

### 2.5. Generic fed-batch fermentation conditions

High cell density fed-batch fermentation was performed at 0.25 L scale (Ambr250 Modular, Sartorius Stedium Biotech).

The MeOH-free fermentation process was initiated at pH 5.0 and 30 °C. Throughout the entire cultivation period, dissolved oxygen concentration was controlled at 30 %. In MeOH-free processes, activity of the PDF promoter is controlled by the addition of a glycerol feed at precisely regulated feeding rates: Fed-batch phase 1 for accumulation of biomass and fed-batch phase 2 for derepression of PDF at a limiting feeding rate. pH and temperature were adjusted for fed-batch phase 2 to support protein production. The total process duration is around 5 days.

### 2.6. Fluorescence measurements

For each strain, samples from three independent cultivations in 24-well deep well plates were measured as biological replicates. Culture samples were diluted in PBS to OD_600_ of 0.5 - 0.6. 100 µL of the OD_600_-adujsted suspension were transferred to a 96-well plate (Corning) and used for the measurement. mNG fluorescence was measured with excitation at 470/15 nm and detection of emission at 515/20 nm, applying a 1000-fold detection gain in a plate reader (CLARIOstar, BMG LABTECH). 10 individual measurements were averaged for each data point. Optical density was measured at 600 nm and values were used to normalize fluorescence signals to biomass.

### 2.7. Total protein quantification in supernatant

As an approximation of the amount of secreted recombinant protein (NANOBODY^®^ VHH B), the total protein in the culture supernatant was determined by the Bradford assay. For each strain, samples from three independent cultivations in 24-well deep well plates were measured as biological replicates. To harvest the supernatant, 24-well deep well plates with cell broth were centrifuged at 4000 rpm for 10 min at 4 °C and 100 µL supernatant was transferred into 96-well plates (Corning). The supernatant was mixed with Quick Start Bradford 1x Dye Reagent (BIO-RAD, 1:1 dilution with MiliQ water). Absorbance of the Bradford samples was measured at 590 nm and 450 nm using a plate reader (Tecan Infinite F200). The Bradford signal was calculated as A_590 nm_/A_450 nm_ (Ernst and Zor, 2010) To convert the Bradford signals into protein concentration, a BSA standard was used.

### 2.8. SDS-PAGE analysis

Cell broth samples were centrifuged (16,000 x g; 2 min) and titers in the cell-free supernatant were quantified using SDS-PAGE/Coomassie staining and software-assisted (Image Lab, BIO-RAD) densitometry scan measurements.

### 2.9. Titer determination by Protein A-HPLC

Titers were determined by Protein A-HPLC as previously described (Groeve et al., 2023).

### 2.10. Documentation of cell floc size

To compare the size of cell flocs, samples from 24-well deep well plates or Ambr250 fed-batch fermentations were diluted to OD_600_ of 5 in complex medium (10 g/L yeast extract, 10 g/L peptone, 1% (w/v) glycerol) and 1 mL of each diluted sample was transferred into a well of a 24-well plate (Corning) and shaken by hand for 10 sec. The plate was imaged using a flatbed scanner (V750 Pro, Epson).

### 2.11. Sedimentation assays

For sedimentation assays, cells were cultured in a 100 mL shake flask filled with 20 mL complex medium (10 g/L yeast extract, 10 g/L peptone, 1 % (w/v) glycerol) at 28 °C for 72 h (80 % humidity, 150 rpm, Kuhner shaker). MeOH-responsive strains were induced by the addition of 1 % MeOH (v/v, final concentration) after 62 h.

For sedimentation assays at natural gravity (1 x g) 10 mL of 72 h-old culture were transferred from shake flask into 15 mL transparent culture tubes (Corning, 16 x 125 mm, polystyrene) and a 50 µL sample of the upper quarter of the suspension was taken to determine OD_600_ using a spectrophotometer (Ultrospec 2100, Amersham Biosciences). After 3 min, another 50 µL of the upper part of the suspension was sampled and optical density was determined. Comparison of the optical density/supernatant turbidity measured at 0 min and 3 min was used to evaluate and quantify the efficiency of sedimentation.

For centrifugation-based sedimentation assays, cell broth samples were diluted in PBS to normalize all samples within each experimental set to an equal OD_600_ value, corresponding to the lowest OD_600_ value in that set. For each cell broth sample, 20 mL of the OD_600_-adjusted suspensions were transferred into 50 mL conical tubes. The conical tubes were centrifuged (Heraeus Multifuge x1r, Thermo Scientific) for 1 min, 2 min, 4 min, and 8 min. From each centrifugation sample, a 50 µL sample of the upper quarter of the suspension was taken at the end of centrifugation to determine the optical density (OD_600_) using a spectrophotometer (Ultrospec 2100, Amersham Biosciences).

Cell sedimentation was additionally photographically documented using a single-lens reflex camera (Canon).

### 2.12. Flocculation inhibition assay

Cells of strain P*_AOX1_*-*ScFLO1* (yEIA004) were cultured in 50 mL shake flasks filled with 20 mL complex medium (10 g/L yeast extract, 10 g/L peptone, 1% (w/v) glycerol) at 28 °C for 72 h (80 % humidity, 150 rpm, Kuhner shaker, ISF1-X). *ScFLO1* expression from P*_AOX1_* was induced by the addition of 1 % MeOH (v/v, final concentration) after 62 h. The culture broth with flocculating cells was adjusted to OD_600_ 30, aliquoted á 2 mL cells were pelleted by centrifugation (3200 rpm, 5 min). Cell pellets were resuspended in either: 2 mL complex medium (10 g/L yeast extract, 10 g/L peptone), 2 mL complex medium with 0.5 M D-glucose, 2 mL complex medium with 0.5 M D-mannose, 2 ml complex medium with 0.05 M D-mannose or 2 mL NANOBODY^®^ VHH A in fermentation medium with a concentration 0.3 g/L and 10 g/L. Floc and pellet formation in different matrices was photographically documented using a single-lens reflex camera (Canon).

### 2.13. Microscopy

Cells suspensions were normalized to OD_600_ 25 in an Eppendorf tube, either by dilution or centrifugation and resuspension. The suspension was afterwards briefly vortexed. After 2 min, 1 µL of cell suspension was taken from the bottom of the Eppendorf tube and transferred onto a glass cover slip. Imaging was performed at room temperature on an optical microscope (DM2500 LED, Leica), equipped with a 40x HCX PL Fluotar objective (NA 0.75) and a digital camera (K3C, Leica). For each sample, single plane images of at least 5 fields of view (corresponding to an average of ∼2000 cells/well) were acquired in brightfield.

## 3. Results

### 3.1. Establishment of a ScFlo1-mediated flocculation phenotype in *K. phaffii*

We first aimed to reproduce the ScFlo1-mediated flocculation phenotype previously described for a *K. phaffii* KM71 strain (Sae-Tang et al., 2023) in a *K. phaffii* NRRL Y-11430 derivative strain. To this end, we intended to site-specifically integrate a single *ScFLO1* expression cassette into the genome to control copy-number and integration locus of the transgene. Targeted, site-specific integration was considered a crucial basis for reproducible characterization of the flocculation phenotype. We expected the *ScFLO1* gene to be a difficult target for classical cloning approaches due to its large size (4614 bp) and the highly repetitive nature of its sequence (Goossens et al., 2011). To avoid an intermediate cloning step, we used CRISPR-assisted targeted integration of the PCR-amplified *ScFLO1* gene into the native *AOX1* locus of *K. phaffii*. This strategy placed *ScFLO1* under control of the methanol (MeOH)-inducible promoter of the endogenous alcohol oxidase 1 gene (P*_AOX1_*). In a second editing step, we replaced P*_AOX1_* with P_GAP_ (Fig. 1b) to enable constitutive *ScFLO1* expression.

For genome manipulations, we used a version of a CRISPR-Cas9 system published for *K. phaffii* (Dalvie et al., 2020), which we modified to be compatible with a method for fast sgRNA cloning (Fig. S1a). In addition, we established a protocol for highly efficient integration of linear dsDNA donors with 80 bp homologies to the target locus (Fig. 1b).

Expression of *ScFLO1* from P_GAP_ in the NRRL Y-11430 derivative strain (yEIA018) led to cell aggregation in complex YPD medium (Fig. S1b) resulting in slightly faster cell sedimentation in a shake flask and culture tube compared to a non-engineered strain. However, we observed dramatic differences in flocculation strength between two YPD batches prepared with peptone from different vendors, with medium 2 not supporting flocculation at all. This medium 2 contained no detectable Ca^2+^ ions (below detection limit) (Fig. S1c), lacking an important ScFlo1-activating cofactor. Addition of CaCl_2_ (ca. 10 mM or 1.1 g/L final concentration) indeed strongly enhanced flocculation in that medium, highlighting the importance of Ca^2+^ ions for efficient flocculation and the careful selection of flocculation supporting growth media.

Encouraged by these results, we sought to further enhance flocculation by increasing *ScFLO1* expression levels. Therefore, we tested flocculation of the strain in which *ScFLO1* is under control of the *AOX1* promoter (P*_AOX1_*) (yEIA010). P*_AOX1_* exhibits 1.5 to 2-fold larger transcriptional activity than P_GAP_ upon respective activating conditions (Vogl et al., 2016). *ScFLO1* expression was induced by MeOH addition after initial carbon source (glucose) depletion. Compared to the P_GAP_-*ScFLO1* strain the floc size increased significantly and became macroscopically visible by eye. Sedimentation in shake flask and culture tube also increased dramatically (Fig. S1b).

### 3.2. Testing flocculation for compatibility with recombinant protein production

Protein glycosylation in *K. phaffii* poses challenges for biopharmaceutical production due to immunogenicity modulation and pharmacokinetic concerns. While N-glycosylation is easily eliminated through protein engineering, O-glycosylation typically manifests as short linear mannose chains (Radoman et al., 2021) with variable frequency and in a product-dependent, unpredictable manner. Since flocculins bind mannose chains in the yeast cell wall to mediate cell-cell adhesion, we investigated whether high concentrations of O-glycosylated recombinant protein in the supernatant could competitively inhibit flocculation.

We performed flocculation inhibition assays by resuspending flocculating cells (P*_AOX1_*-*ScFLO1,* yEIA004 induced with 1 % MeOH) in media containing free D-mannose and D-glucose or a representatively O-glycosylated pentavalent NANOBODY^®^ VHH molecule. The degree of O-glycosylation of NANOBODY^®^ VHH A was determined by mass spectrometry to be 19 % to 20 % (single O-hexosylation (1x): 11%, 2x: 2.5%, 3x: 0.7%, 4x: 0.3%). The positive control treatment (0.5 M D-mannose) inhibited flocculation quantitatively, while adding D-glucose (negative control) at equivalent concentration showed no effect. These results are in line with previous characterization of ScFlo1 competitive binding inhibition (Degreif et al., 2017). Notably, no inhibition of flocculation was observed in the presence of 10 g/L glycosylated NANOBODY^®^ VHH A corresponding to approximately 0.028 mM to free mannose equivalents (Fig. 1e).

To test our flocculation system under conditions closer to biopharmaceutical production, we genomically integrated NANOBODY^®^ VHH B expression cassettes into the strains already equipped with the *ScFLO1* expression cassette. The NANOBODY^®^ VHH B expression cassette consisted of the PDF promoter, and a sequence encoding the full-length alpha mating factor secretion leader of *S. cerevisiae* fused in-frame with the NANOBODY^®^ VHH B encoding sequence (Groeve et al., 2023). The derepressible PDF promoter is a 623 bp fragment of the formate dehydrogenase promoter of *Hansenula polymorpha* that mediates high-level expression under carbon source limiting conditions in *K. phaffii*. It can additionally be activated by methanol (Vogl et al., 2020), allowing full compatibility with *ScFLO1* expression from either P_GAP_ (yIEA038) or P*_AOX1_* (pEIA024). Two expression cassettes were integrated into each strain at loci on chromosome 2 (ChrII:17928–17947 = int6) and chromosome 4 (ChrIV:733170–733189 = int18) described in (Gao et al., 2022) using the CRISPR-Cas9 system. In accordance with the results described above, the degree of cell-cell aggregation in these strains was significantly stronger for *ScFLO1* expression from P*_AOX1_* than from P_GAP_ (Fig. 1c). Consistently, P*_AOX1_*-*ScFLO1* cells exhibited enhanced sedimentation rates: 3 min after transferring the cell suspension to culture tubes, these cells achieved near-complete separation from the supernatant (Fig. 1d). A non-flocculating reference strain in which *ScFLO1* was replaced by the fluorescent reporter mNeonGreen (mNG) (P*_AOX1_*: yEIA025) showed no observable sedimentation during the equivalent time period. Cells expressing *ScFLO1* from P_GAP_ did not sediment markedly faster than the respective mNG-expressing reference cells (P_GAP_: yIEA039) (Fig. 1d). Expression assays with these strains in 96-well deep well plates demonstrated that mNG reference strains and flocculating strains produced similar level of NANOBODY^®^ VHH B (Fig. S2a, b & Fig. S3). Hence, flocculation does not have any negative impact on strain productivity.

### 3.3. Advancing inducible flocculation phenotype with genetic amplifiers

Compared to the flocculation we observed so far, previous studies in other yeast species had demonstrated significantly stronger flocculation upon *ScFLO1* overexpression (Degreif et al., 2017; Smukalla et al., 2008). This prompted us to investigate if flocculation can be enhanced in our P*_AOX1_*-*ScFLO1* strains by further increasing the expression of *ScFLO1*.

To this end, we genomically integrated additional genetic amplifier modules in the genome containing either *MIT1* (*ScFLO1*: yEIA042, mNG: yEIA044) or *MXR1* (*ScFLO1*: yEIA043, mNG: yEIA045), both under P*_AOX1_* regulation. Both designs create synthetic positive feedback circuits (Fig. 2a), as the transcriptional activators (TA) Mit1 and Mxr1 upregulate the P*_AOX1_* promoter (Lin-Cereghino et al., 2006; Wang et al., 2016; Zhan et al., 2017) that drives their own expression and simultaneously activate P*_AOX1_*-driven *ScFLO1* expression. Such genetic amplifiers have been shown to significantly boost recombinant protein expression levels upon MeOH induction (Poodeh et al., 2022).

**Fig. 2:**
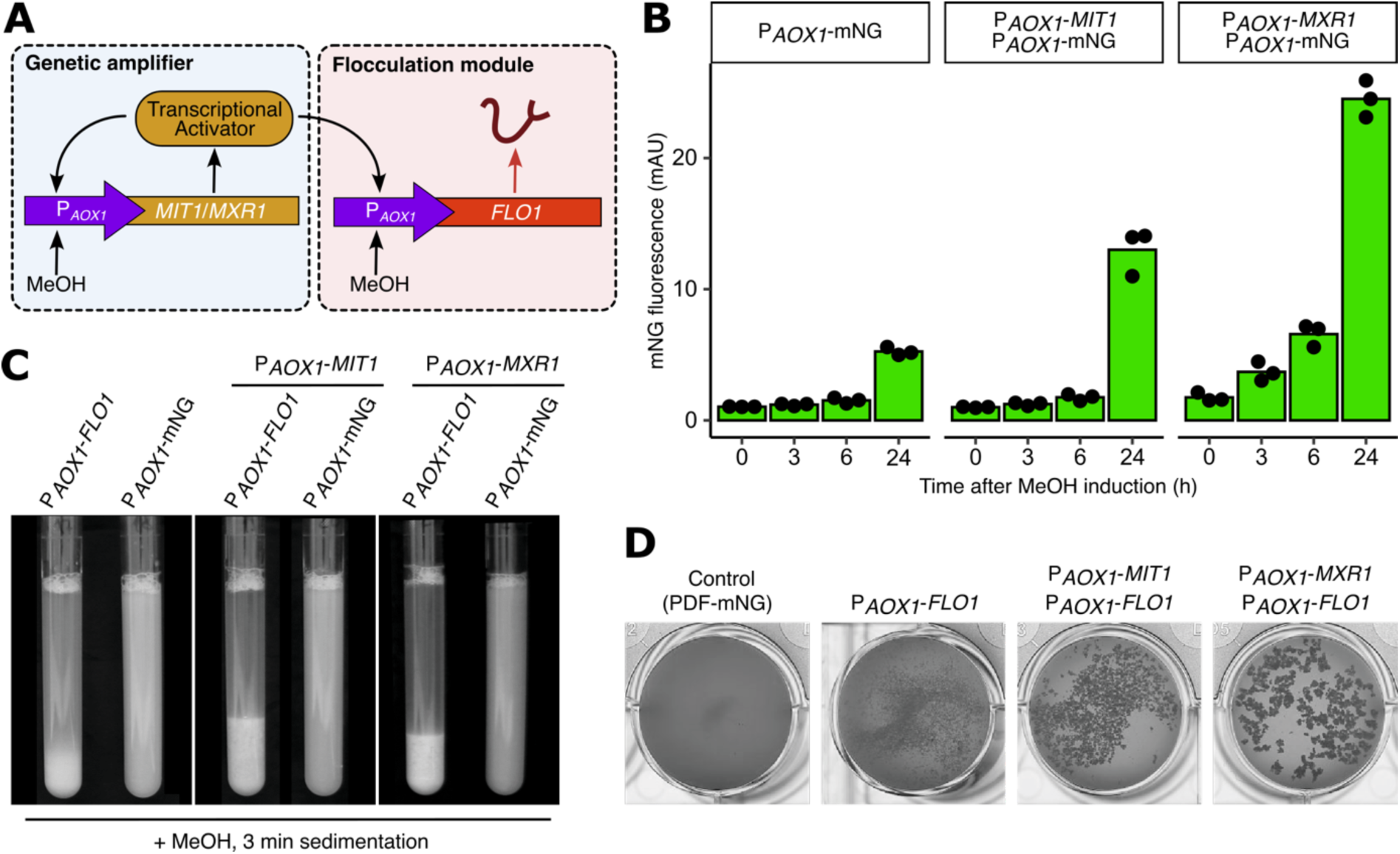
Enhancing inducible flocculation with genetic amplifiers. **A:** Schematic representation of the genetic amplifier module: Transcription activators Mit1 and Mxr1 are under control of P*_AOX1_* which they are activating thus creating a synthetic positive feedback loop. Transcription activators Mit1 and Mxr1 are additionally activating *ScFLO1* expression from P*_AOX1_*. **B:** Fluorescence levels upon mNeonGreen (mNG) expression from P*_AOX1_* in the absence or presence of genetic amplifier modules to measure MeOH induction kinetics (0 h, 3 h, 6 h, 24 h) and absolute expression strength. n = 3 **C:** Sedimentation behavior of indicated strains after growth in shake flask and MeOH induction. Cells were transferred to culture tubes and photographed after 3 min. **D:** Microscopic comparison of the floc size of indicated strains. For macroscopic analysis, cell suspension was transferred into a well of a 24-well plate.

To assess the impact of genetic amplifier constructs on P*_AOX1_*-mediated expression, we constructed reporter strains in which mNG expression is driven by the *AOX1* promoter. Indeed, MeOH-induced overexpression of either activator led to much higher expression levels of mNG upon methanol induction. *MXR1* co-expression mediated the strongest fluorescence level at all time points (Fig. 2b). 24 h after MeOH-induction mNG expression levels were about 2.5-fold higher with *MIT1* co-expression and 5-fold higher with *MXR1* co-expression than without genetic amplifier. Accordingly, increased floc size was observed for both P*_AOX1_*-*ScFLO1* strains containing either genetic amplifier. *MXR1* co-expression increased flocculation so drastically, that large flocs became visible by eye (Fig. 2d). Large floc size improved sedimentation speed and led to clearer supernatant after 3 min sedimentation in a culture tube (Fig. 2c, Fig. 5b). NANOBODY^®^ VHH B expression levels were not affected by the *MXR1*-enhanced flocculation phenotype (Fig. S2b) and secretome and product quality remained unaffected, as determined by SDS-PAGE of the culture supernatant (Fig. S3). Based on these results, we identified the *MXR1*-based genetic amplifier as advantageous for our purposes.

### 3.4. Development of a MeOH-independent inducible flocculation system

Eliminating methanol (MeOH) from industrial production processes offers substantial advantages due to MeOH’s flammability and toxicity. We, therefore, aimed to develop a flocculation system inducible through carbon source limitation, leveraging the PDF promoter. Accordingly, we wanted to determine whether the PDF promoter could drive expression to levels sufficient to meet the requirements of our flocculation system. To evaluate this, we replaced the P*_AOX1_* promoter driving mNG expression with the PDF promoter in strains already containing a PDF-driven NANOBODY^®^ VHH B expression cassette, enabling co-expression of VHH and mNG upon carbon source limitation.

This reference strain expressing mNG from PDF (yIEA053) demonstrated significantly higher expression levels following carbon limitation-induced activation compared to P_GAP_-mNG (yIEA039) and P*_AOX1_*-mNG (yIEA025) after methanol induction (Fig. 1c & Fig. 3c). These observations led us to hypothesize that employing the PDF promoter might enhance flocculation efficiency beyond previously observed levels in strains lacking a genetic amplifier.

**Fig. 3:**
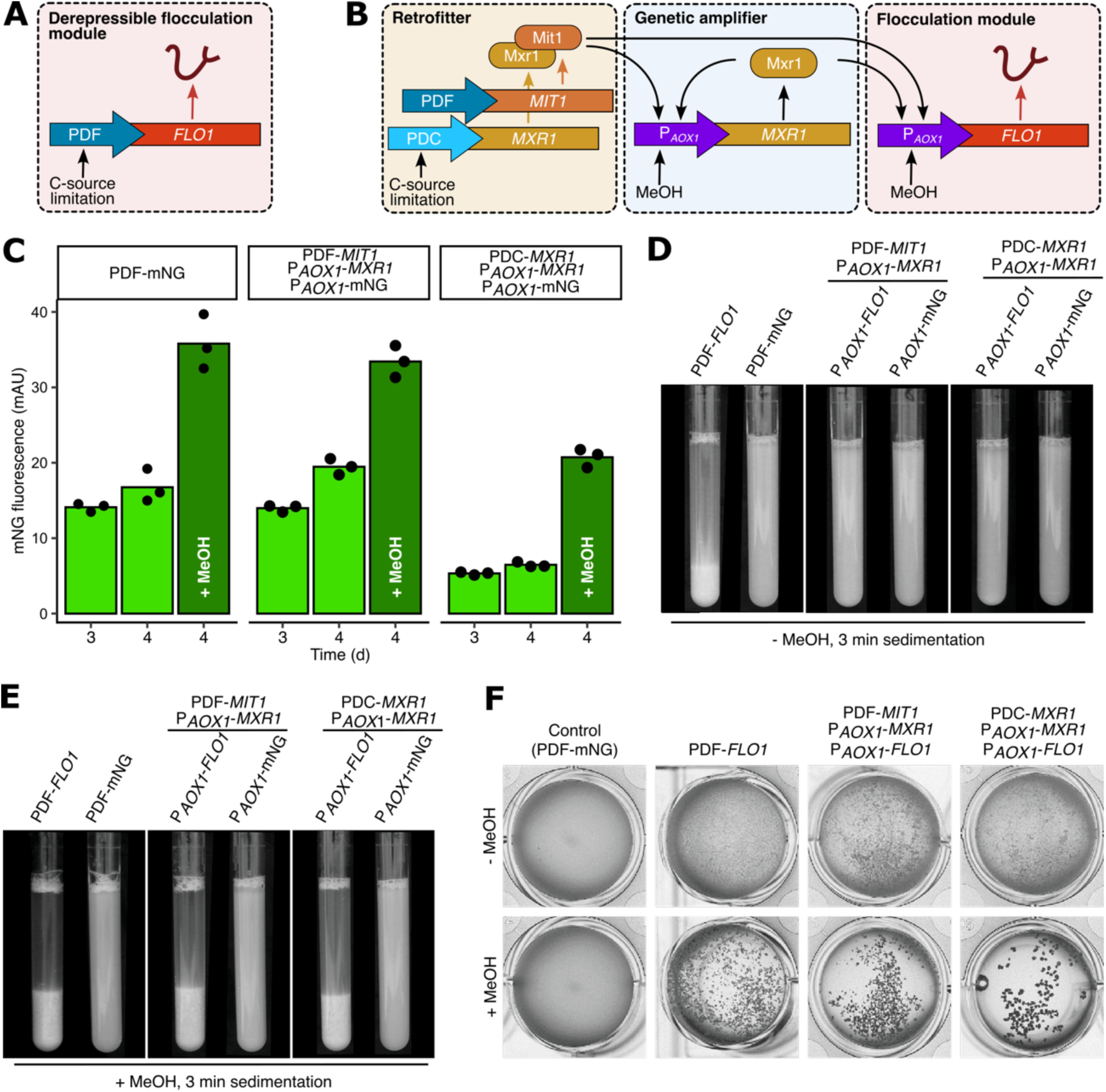
Development of a MeOH-independent inducible flocculation. **A:** Schematic representation of the PDF-*ScFLO1* flocculation module allowing for activation upon derepressing conditions at limited carbon source concentrations. **B:** Schematic representation of the interaction of the retrofitter module with the genetic amplifier module and the P*_AOX1_*-based flocculation module. Derepressing conditions at limited carbon source concentrations induce *MXR1*/*MIT1* expression from the retrofitter, which then activates the genetic amplifier module. **C:** mNG fluorescence levels upon expression from the PDF promoter or P*_AOX1_* in the presence of the genetic amplifier module (P*_AOX1_*-*MXR1*) and either version of the retrofitter module (PDF-*MIT1* or PDC-*MXR1*) after 3 and 4 days and after induction with MeOH. n = 3 **D:** Sedimentation behavior of indicated MeOH-independent strains after growth in shake flask without MeOH induction. Cells were transferred to culture tubes and photographed after 3 min. **E:** Sedimentation behavior of identical strains as analyzed in D, with MeOH-induction to test the additional MeOH-activation of the PDF promoter in comparison to the fully induced and P*_AOX1_*-*MXR1* amplified P*_AOX1_*-*ScFLO1* strains. Cells were transferred to culture tubes and photographed after 3 min. **F:** Macroscopic comparison of the floc size of indicated strains without and with MeOH-induction. As non-flocculating reference, a PDF-mNG strain is included. For macroscopic analysis, cell suspension was transferred into a well of a 24-well plate.

To test this hypothesis, we changed the P*_AOX1_* of *ScFLO1* in strain yEIA025 to the PDF promoter (Fig. 3a), so that again both NANOBODY^®^ VHH B and *ScFLO1* are co-expressed upon carbon source limitation. As expected, the PDF-Sc*FLO1* construct (yIEA052) showed a strong flocculation phenotype upon promoter derepression (Fig. 3d). Turbidity of the supernatant after cell sedimentation for 3 min (Fig. 3d) and sedimentation speed (Fig. 5b) were similar to previous observations for the genetic amplifier constructs induced with MeOH. In line with the previous results, PDF-driven *ScFLO1* co-expression did not affect NANOBODY^®^ VHH B expression (Fig. S2b & Fig. S3). In addition to derepression upon C-source limitation, the PDF promoter can be further activated by MeOH (Vogl et al., 2020). To test if this characteristic of the PDF promoter could be exploited, we also induced the PDF-based co-expression strain variants using MeOH. Fluorescence intensity for PDF-mNG increased more than two-fold upon MeOH addition (Fig. 3c). In accordance, floc size (Fig. 3f) and sedimentation speed (Fig. 5b) also increased upon MeOH induction of the PDF-*ScFLO1* strain leading to a clearer culture supernatant after 3 min cell sedimentation (Fig. 3e).

As both genetic amplifier strains exhibited enhanced floc size following MeOH induction compared to the PDF-*ScFLO1* construct in MeOH-free conditions, we investigated whether a gene circuit could be engineered to replicate this strong flocculation phenotype independently of induction by MeOH. To achieve MeOH-independent flocculation, we employed retrofitter modules (Fig. 3b), in which the P*_AOX1_*-activating transcription factors Mxr1 or Mit1 are placed under the control of derepressible promoters (Vogl et al., 2018). Based on our previous findings, we designed two complementary regulatory configurations to balance induction sensitivity and transcriptional leakiness of *ScFLO1* expression. First, we paired the stronger PDF promoter with the weaker activating factor Mit1 (*ScFLO1*: yEIA047, mNG: yEIA048). Second, we combined the weaker derepressible PDC promoter (CAT1-100 promoter from *K. phaffii* (Fischer et al., 2019) with the stronger activating factor Mxr1 (*ScFLO1*: yEIA049, mNG: yEIA050). In these engineered strains, carbon-source limitation triggers the expression of Mxr1 or Mit1, respectively. These transcription factors then activate *ScFLO1* expression from P*_AOX1_*. Simultaneously, they induce the self-amplifying P*_AOX1_*-*MXR1* module, which further enhances and maintains P*_AOX1_*-*ScFLO1* expression, creating a positive feedback loop that amplifies the flocculation response (Fig. 3b).

The flocculation phenotype of both constructs exhibited similar strength compared to the MeOH-independent PDF-*ScFLO1* strain, as evidenced by comparable floc sizes (Fig. 3f). The increased floc size consequently translated into substantially enhanced sedimentation rates (Fig. 5b) although these were slightly reduced relative to PDF-*ScFLO1*. The sedimentation of these retrofitted strains also appeared visually weaker in culture tube (Fig. 3d).

Under MeOH-free conditions, the mNG reference strain containing the PDF-*MIT1* module (yEIA048) demonstrated slightly higher fluorescence levels compared to its PDF-mNG reference counterpart. Conversely, PDC-*MXR1* (yEIA050) exhibited fluorescence levels (Fig. 3c) even lower than previously observed for P_GAP_-mNG (Fig. 1c). The addition of MeOH resulted in strongly enhanced mNG fluorescence expression levels for both retrofitter constructs (Fig. 3c) as well as increased floc size (Fig. 3f). While these changes led to improved sedimentation speed (Fig. 5b), the supernatant turbidity after 3 min of cell sedimentation remained slightly higher than that observed for PDF-*ScFLO1* under equivalent MeOH conditions (Fig. 3e).

NANOBODY^®^ VHH B expression levels showed construct-dependent responses. Expression levels remained stable in strains overexpressing *MXR1* from PDC but decreased significantly in strains overexpressing *MIT1* from PDF (Fig. S2b & Fig. S3).

### 3.5. Testing flocculation in bioreactor conditions

In the previously described experiments, we have characterized the flocculation phenotype of our engineered strains by using cells that were cultured in 96-well deep well plate format or in a shake flask model (for sedimentation assays only). As we aimed to investigate impact of genetically engineered cell aggregation for supernatant-biomass separation in industrial, cubic meter scale production processes, we next needed to test for compatibility of our engineered flocculation strains with more representative cultivation conditions in a bioreactor system. In contrast to shake flask cultivation, fermentation in a bioreactor allows for improved control of pH, dissolved oxygen and a feeding regime to finely adjust carbon source availability. Such substantial changes in culture conditions can impact flocculation factors such as cell density, *ScFLO1* expression and cell wall composition including relative amounts of mannans (Farinha et al., 2022). Furthermore, cells are simultaneously exposed to harsher conditions like intense shear stress, not only during fermentation but additionally during harvest operations.

Fully committed to MeOH-free fermentation processes to prioritize operator safety, facility protection, and environmental sustainability, we selected the PDF-*ScFLO1* (yIEA052) construct *-* the MeOH-independent strain variant that demonstrated strongest flocculation in our small-scale characterization studies - for evaluation in a bioreactor system (Ambr250; Sartorius AG). The corresponding weaker MeOH-independent flocculant strain P_GAP_-*ScFLO1* was included as a reference. Both strains are fully compatible with established MeOH-free fermentation (Groeve et al., 2023). The flocculating strains were additionally equipped with two copies of the PDF-based NANOBODY^®^ VHH B expression cassette as described above. Respective mNG reference strains served as controls, while a strain version containing only two copies of the NANOBODY^®^ VHH B expression cassette without *ScFLO1* integration (yIEA056) was used to compare productivity.

Both the P_GAP_-*ScFLO1* and PDF-*ScFLO1* strains achieved cell densities similar to all control strains (approximately OD_600_ of 400, Fig. S2c) although the PDF-*ScFLO1* strain exhibited a slightly reduced growth rate relative to the P_GAP_-*ScFLO1* and the control strains as deduced from measured oxygen uptake rates (OUR) (Fig. 4a). Flocculation occurred in both *ScFLO1* expressing strains at the end of the fed-batch phase. Floc formation was detectable by microscopy and was macroscopically visible in culture broth transferred to 24-well plates (Fig. 4c). Despite macroscopic floc formation, no differences in sedimentation kinetics were observed under natural gravitation (1 x g). We attributed this observation to the high viscosity of the cell broth. To simulate industrial-scale biomass separation using a disc-stack separator (continuous centrifugation), culture broth aliquots were subjected to a centrifugation-based sedimentation assay (> 1 x g, Fig 5a). Culture broths from different strains were normalized to OD_600_ 350, and 20 mL aliquots were centrifuged at low speed (1000 x g) for durations ranging from 1 min to 8 min. Optical density measurements of the resulting supernatants revealed substantial differences in sedimentation behavior among strains, with the PDF-*ScFLO1* strain sedimenting most rapidly, followed by the P_GAP_-*ScFLO1* strain, which sedimented faster than both the non-flocculating strain (yIEA056) and the respective mNG control strains (Fig. 5c). This trend correlated directly with flocculation strength (Fig. 4c). After only 1 min of centrifugation, the supernatant of the PDF-*ScFLO1* strain culture exhibited three-fold lower turbidity compared to non-flocculating strain variants (approximately OD_600_ = 10 vs. OD_600_ = 30). Significant differences in supernatant optical density persisted through the 4 min time point (Fig. 5c, d). After 8 min of centrifugation, maximal clarification was achieved at the applied g-force, and no further differences were detectable (Fig. 5c). Productivity, as determined by recombinant protein concentration in the supernatant, remained identical across all strains (Fig. 4b). Product quality, as assessed through a comprehensive analytical panel (Tab. S6), remained unchanged by the induction of the flocculation phenotype. Where significant relative differences were observed compared to the reference strain expressing NANOBODY^®^ VHH B alone, these variations were not exclusive to the *ScFLO1*-expressing strains but were concurrently detected in corresponding mNG reference strains. Consequently, such differences are attributable to the co-expression of additional heterologous proteins (*Sc*Flo1p or mNG) rather than to the flocculation phenotype *per se*.

**Fig. 4:**
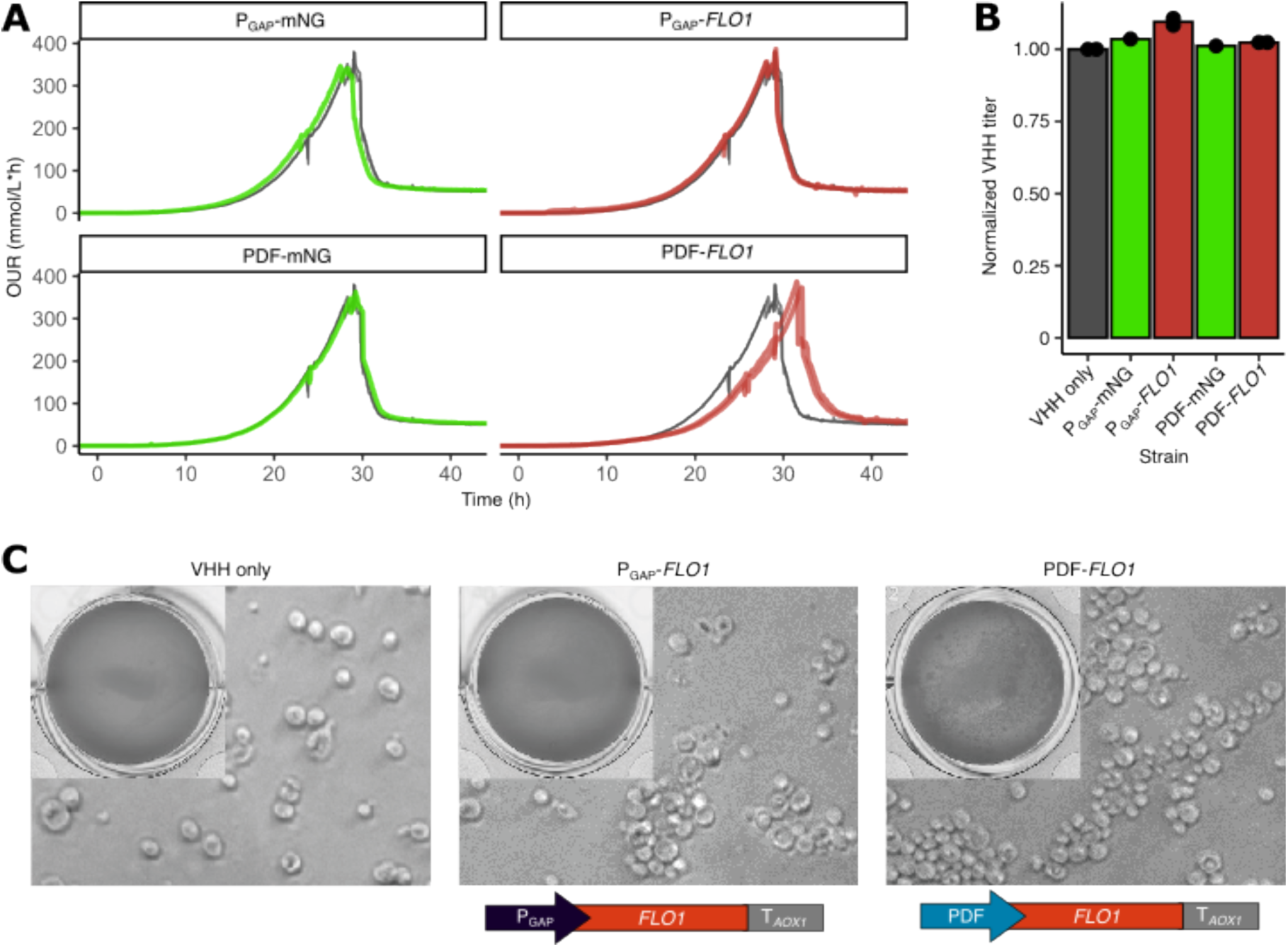
NANOBODY^®^ VHH B production in flocculating strains. **A:** Growth evaluation of indicated strains in Ambr250 bioreactor cultivations, as measured by oxygen uptake rate (OUR). The empty strain reference expresses NANOBODY^®^ VHH B only and does not harbor a *ScFLO1* expression cassette. Each strain was cultured in two independent technical replicates. **B:** Comparison of productivity of indicated flocculating strains and control strains in Ambr250 bioreactor cultivations. Productivity (product titers) are normalized to the empty reference strain expressing NANOBODY^®^ VHH B only. **C:** Microscopic and macroscopic comparison of the floc size of indicated strains from Ambr250 bioreactor cultivations. For macroscopic analysis, cell suspension was transferred into a well of a 24-well plate (top left corner). Scale bar = 10 µm.

**Fig. 5:**
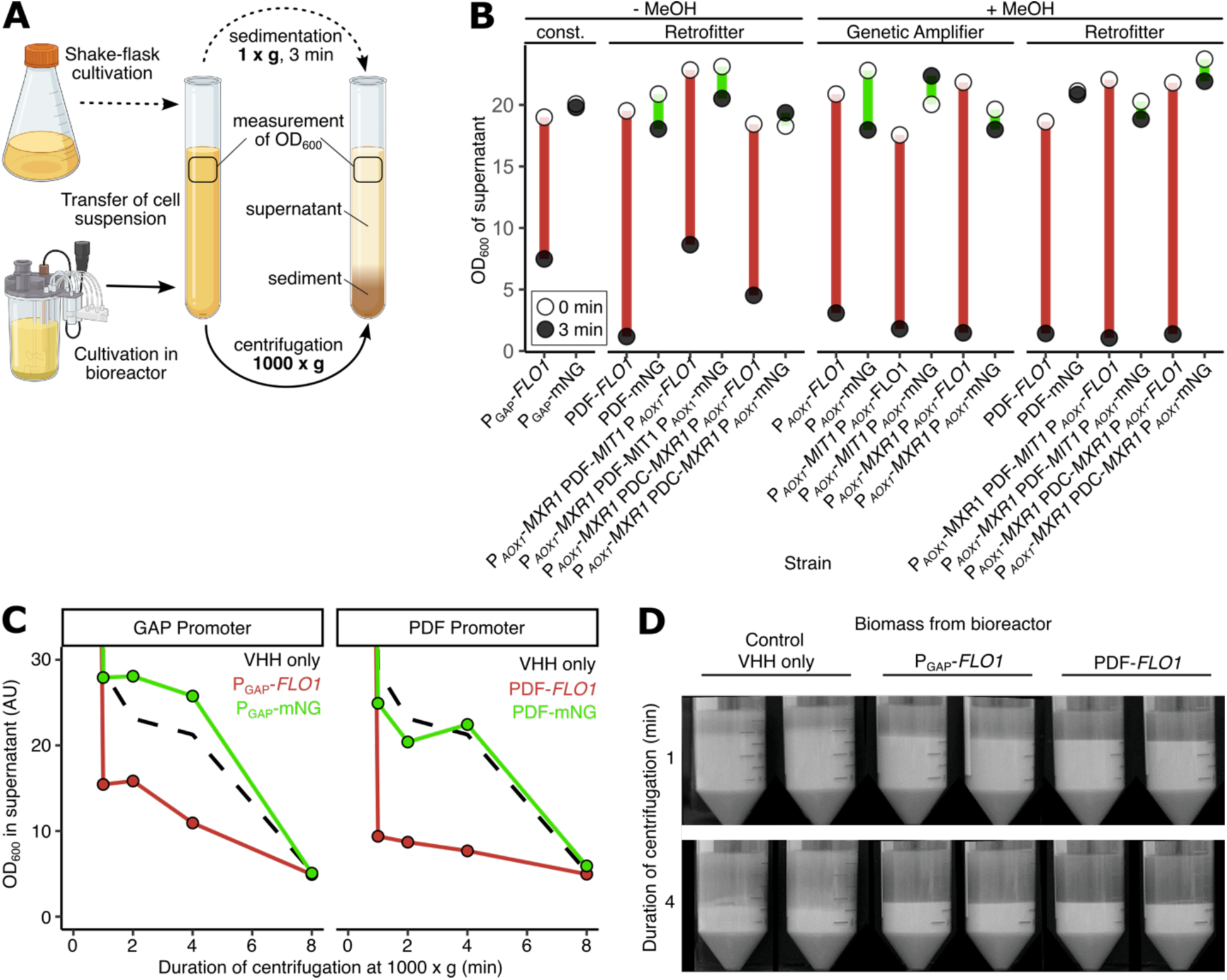
Analyses of sedimentation speed and efficiency. **A:** Schematic representation of sedimentation analysis. Cell suspensions from shake flask or Ambr250 bioreactor cultures were transferred to 15 mL tubes (sedimentation) or 50 mL conical tubes (centrifugation) and allowed to sediment under natural gravity (1 x g) for 3 min or centrifuged (> 1 x g) for 1, 2, 4, or 8 min. A 50 µL sample was respectively collected from the upper quarter of each tube, and OD_600_ was measured and compared to the initial OD_600_ of the transferred cell suspension. Created in BioRender. Degreif, D. (2026) https://BioRender.com/54nbqy4 **B:** Sedimentation speed analysis of indicated strains grown in shake flask under specified conditions (-MeOH, +MeOH) as indicated in A. **C:** Sedimentation speed analysis of indicated strains grown in Ambr250 bioreactor. Individual samples were centrifuged (1000 x g) for 1, 2, 4 or 8 min respectively and analysis was performed as indicated in A. In black (dashed lines), identical analysis of a strain expressing NANOBODY^®^ VHH B only (yEIA056) is shown. **D:** Representative exemplary images of the optical differences in separation efficiency upon centrifugation at 1000 x g for 1 min and 4 min for indicated strains cultured in Ambr250 bioreactor. The samples correspond to the analyses shown in C.

## 4. Discussion

Most studies to improve recombinant expression hosts focus on increasing productivity or quality of the product. Yet, in an industrial setting, high protein yields are only part of the prerequisites for efficient manufacturing processes. *K. phaffii* enables high protein productivity, shifting efficiency bottlenecks to process robustness and downstream yield. In particular, separating large quantities of dense biomass from product-containing supernatant has only been addressed insufficiently until now.

In this study, we engineered *K. phaffii* strains to flocculate upon induction and demonstrate, for the first time, the combination of genetically engineered cell aggregation with recombinant protein expression to facilitate separation of product and biomass. We successfully demonstrated the general feasibility of this approach using biomass generated in controlled bioreactor cultivations, showcasing that enhanced biomass separation under both natural gravity and centrifugation conditions can be achieved without compromising productivity or product quality. This was validated by expressing a complex pharmaceutically relevant protein, confirming the applicability of our technology to industrial biopharmaceutical manufacturing. While flocculation is well-established in brewing industry and wastewater treatment, its integration into biopharmaceutical manufacturing faces stricter regulatory barriers and demanding quality requirements. Chemical flocculants introduce additional expensive raw materials that must meet stringent pharmaceutical quality standards. Relying on environmental triggers to induce wildtype flocculation pathways typically exploited in brewing industry such as nitrogen starvation, hypoxic conditions occurring towards to end of fermentation or the presence of cytotoxic compounds such as ethanol is also not compatible with well-controlled fermentation processes applied in biopharmaceutical industry.

Therefore, we aimed to establish flocculation as an engineered, inducible strain feature eliminating the need for external flocculants or triggering by environmental stressors. Genetic regulation additionally enables precise temporal control of flocculation initiation. This temporal control is considered crucial since early floc formation can limit nutrient diffusion to cells inside the flocs, thus reducing growth, biomass production and potentially impacting product formation and quality. While such diffusion barriers naturally protect wildtype yeast from environmental threats (Smukalla et al., 2008), they are undesirable in well-controlled production settings. Therefore, flocculation must be activated late in the fermentation process after sufficient biomass was accumulated. In accordance, we successfully engineered inducible flocculation systems in which derepressible promoters simultaneously control both, product expression and the initiation of flocculation. Lacking strong and tight promoters for *K. phaffii* that can be induced by non-toxic and non-flammable compounds, we placed the flocculin under P*_AOX1_* control to activate flocculation orthogonally in an alternative system. Both approaches allow to induce flocculation in fermentation when biomass accumulation is complete, and cell growth is minimal. The implementation of genetic amplifiers significantly enhanced the P*_AOX1_*-mediated flocculation. Additional amplifier configurations, such as the co-expression of *MXR1* and *MIT1*, which were not evaluated in this study and may be subject of future studies, could even further intensify flocculation strength. However, P*_AOX1_*-controlled flocculation strains require explosion-proof facilities. Therefore, the MeOH-free systems that we have engineered offer even broader compatibility with existing biopharmaceutical production facilities, enhance operator and facility safety, and better comply with environmental requirements. Nevertheless, MeOH-induction provides advantages through expression control independent of product formation. Future work will address orthogonal induction of product and flocculation using safe, environmentally friendly and industry-compatible methods. Also, the retrofitting concept evaluated in this study presents significant potential for further optimization. Future investigations could e.g. explore alternative promoter-transcription factor combinations, such as pairing the more potent PDF promoter with the enhanced activating factor Mxr1. In general, our study primarily examined variations of promoter systems to optimize *FLO* gene transcription. However, we recognize significant potential for further improvement of our system in several areas: translation efficiency, protein processing in the secretory pathway, protein turnover and the mount of active protein at the membrane. Additionally, exploring flocculins derived from alternative yeast species or engineered chimeric variants presents promising avenues for future research. Overall, our lab scale results demonstrate that the genetic amplifier and retrofitter concepts can be used to regulate additional functional features in production strains and are not limited to boost and control product expression. In particular, the transcription cascade introduced by retrofitting circuits introduces a temporal delay which might be exploited when sequential expression of product and auxiliary proteins is beneficial.

P_GAP_-mediated expression is well-known to be strongly coupled with growth, exhibiting maximal promoter activity at high growth rates (Garcia-Ortega et al., 2016; Nieto-Taype et al., 2020). Consequently, in a previously published system where P_GAP_ controls *ScFLO1* expression, flocculation is initiated when cells are actively growing, which is an issue in an industrial protein production setting, as discussed above. We therefore considered decoupling of cell growth and flocculation initiation to be crucial for preventing any impairment of biomass formation by pre-mature flocculation. Surprisingly, our control experiments using P_GAP_-*ScFLO1* constructs revealed no growth-inhibiting effects, even during bioreactor cultivations that generated substantial biomass quantities. However, P_GAP_-mediated flocculation presented in literature (Sae-Tang et al., 2023) had been described to be stronger than the phenotype observed in this study. These differences cannot be attributed to differences in strain genetic background, since *K. phaffii* strains KM71 and NRRL Y-11430 are closely related: KM71 is an mut^S^ variant of GS115 (Karbalaei et al., 2020) which is a *his4* auxotrophic mutant of NRRL Y-11430 (Claes et al., 2024). In our engineered strains, integrations into the *AOX1* locus even created a mut^S^ phenotype so that the only difference is the *his4* mutation. Instead, differences in *ScFLO1* copy number and the integration loci are more likely to contribute to the observed differences in flocculation strength. The P_GAP_-*ScFLO1* KM71 strain was generated by random integration of an integrative plasmid with no information on the copy number and integration locus. Random integration often results in multi copy strains, which aligns with the stronger flocculation phenotype reported for the KM71 strain-based study. However, we consider the typically concatemeric architecture of randomly integrated vectors (tandem integrations) (Schwarzhans et al., 2016) less favorable for industrial applications due to elevated risks of genetic instability by loop-out recombination. Our findings demonstrate that the inducible expression systems we developed exhibit stronger flocculation compared to P_GAP_-mediated flocculation on a per *ScFLO1*-copy basis. This enhanced flocculation performance at low copy number in combination with their inducible character renders them particularly advantageous for industrial applications, beyond genetic stability considerations. We hypothesize that stronger P_GAP_-mediated flocculation from multi-copy strains would likely severely impair cell growth by pre-mature flocculation. In line with this hypothesis, the minor growth impairment in the PDF-*ScFLO1* strain (yIEA052) also likely results from weak pre-mature flocculation assuming that the PDF promoter is already partially active during the feeding phase of the fermentation process. Since PDF is 4- to 5- times stronger than P_GAP_, even partial PDF activation could cause flocculation stronger than the fully activated P_GAP_ mediates at single *ScFLO1* copy levels. However, this minor growth defect is negligible given the ability to achieve identical final biomass and the strong flocculation phenotype at end of fermentation. These results rather underscore the critical need for temporal separation between biomass formation and flocculation, emphasizing the importance of developing strong flocculation systems with even better temporal control than already achieved with PDF-*ScFLO1* (yIEA052).

However, flocculin expression levels and timing represent just one contributing factor to flocculation strength. Equal attention must be paid to media and matrix composition, as the mannose-binding activity of ScFlo1p exhibits significant sensitivity to both pH conditions (Jin and Speers, 2000; Soares, 2011) and divalent ion (especially Ca^2+^) concentrations (Fig. S1c) (Miki et al., 1982). Therefore, defined synthetic media are preferred over media with complex components (such as hydrolysates) as they allow for full control of media composition. Defined media also allow for control of flocculation by tuning Ca^2+^ concentration during fermentation. Other environmental factors such as temperature and pH must be tightly controlled during the fermentation process to ensure optimal flocculation, thus introducing additional technical challenges.

The successful industrial implementation of engineered expression strains with enhanced processability characteristics requires consideration of multiple interdependent factors beyond the inherent strain performance. Critical determinants include compatibility with established, regulatory-approved production processes and integration with existing technical infrastructure within GMP-compliant manufacturing environments. When strain-based processability improvements necessitate substantial modifications to production facilities, manufacturers face significant capital investments and operational downtime during facility upgrades. The economic advantages gained from enhanced processability must therefore demonstrate sufficient return on investment to offset these substantial costs, creating a significant commercial barrier to novel strain adoption in industrial settings. Furthermore, the complexity of associated regulatory compliance requirements presents additional implementation challenges. Our findings represent a promising initial advancement toward incorporating engineered flocculation into routine commercial production workflows. Beyond large-scale manufacturing applications, this technology demonstrates considerable potential for small-scale laboratory implementations, particularly as a streamlined biomass separation method in high-throughput screening protocols for recombinant protein expression studies.

The cell flocculation technology presented in this study demonstrates compatibility with established harvesting operations utilizing continuous disc-stack centrifugation followed by depth filtration, as well as with alternative sedimentation tank approaches, which, although uncommon in pharmaceutical applications, could be considered for new facilities or facility remodeling projects. When cell flocs are processed through commonly used continuous disc-stack centrifugation, we expect biomass separation to become more efficient even without changing the gravitational field strength or the average biomass residence time within the disc-stack centrifuge. This improvement follows the physical principles described by Stokes’ law with cell aggregates creating larger particles that separate more effectively under unmodified centrifugation conditions, thus reducing the biomass leaking to the centrate. Cell flocculation could offer an additional benefit to improve centrate clarity by allowing the flocculated biomass to bind and capture small cell fragments and cellular membrane debris from lysed cells, removing particles that would otherwise remain in suspension due to their small size. A clearer centrate would then reduce the number of depth filtering steps with considerable impact on process time and_costs. However, given the lack of reliable small-scale models for disc-stack separation and depth filtration processes, these theoretically derived advantages necessitate experimental validation through pilot-scale studies to confirm their practical applicability. Further fermentation process development is essential to establish complete compatibility between genetically engineered flocculation and bioreactor-based fermentation processes. While our proof-of-principle studies demonstrate general feasibility, scaling beyond laboratory conditions requires optimization to maintain the robust flocculation phenotypes observed during small-scale expression studies. Additionally, process development efforts must systematically address the impact of strain engineering modifications on critical product quality attributes, as evidenced by the bioreactor cultivation results presented (Tab. S6).

## 5. Acknowledgments

We thank Marcel Boss, Robert Gnügge, Philipp Höß, Melanie Knoll, Alexander Leske, Dorine Phan, Louisa Rothenberger, Kathleen Schneider, Nicole Schottstedt, Jessica Swiridow and Sylvia Zink for valuable comments on the manuscript.

This work was fully funded by Sanofi.

## 6. Author contributions

E.I, P.R., N.Z. performed experiments and analyzed data. E.I., M.M., E.A., C.S. and D.D. conceived the study. M.M., C.S. and D.D. supervised the study. E.I., C.S. and D.D. wrote the manuscript with input from the other authors.

