## Supplemental Information for "CRISPR-engineered inducible flocculation in *Komagataella phaffii* enables enhanced biomass separation for biopharmaceutical production"

This file contains 6 tables and 3 figures.

### Tab. S1: Oligonucleotides used in this study

| Oligo | Sequence (5' → 3') |
|---|---|
| O1 | TTCCAATTGACAAGCTTTTGATTTTAACGACTTTTAACGACAACTTGAGAAGATCAAAAAACAACTAATTATT CGAAACGATGACAATGCCTCATCGC |
| O2 | CGATAAGACCAACTTTCAAGGAGTGGTCCAAGTTTGCCAATCTTCCGGCAATACAGGATCCACTGGATCC ACCATTAAATAATTGCCAGCAATAAGG |
| O3 | GGTGGGAATACTGCTGATAGCC |
| O4 | ACCATCTGGAAGTGTCACAC |
| O5 | TTAGCGGCGTCACAACAG |
| O6 | CCTGGAAGGTAGACCCATG |
| O7 | TTCCAATTGACAAGCTTTTGATTTTAACGACTTTTAACGACAACTTGAGAAGATCAAAAAACAACTAATTATT CGAAACGATGGTTTCTAAGGGTGAAGAGG |
| O8 | CGATAAGACCAACTTTCAAGGAGTGGTCCAAGTTTGCCAATCTTCCGGCAATACAGGATCCACTGGATCC ACCATTACTTGTACAACTCGTCCATACC |
| O9 | GATGATTATGCATTGTCTCCAC |
| O10 | GTTGGTATTGTGAAATAGACGCAGATCGGGAACACTGAAAAATAACAGTTATTATTCGAGATCTAACATCC AAAGACGAATTTTTGTAGAAATGTCTTGGTGTCC |
| O11 | CCGTTGATGGAACCGAAAATGTGCAACTCGTGAGTAGCTGGCAAGGAAGCCATGTTGTCCTCTTCACCCT TAGAAACCATTGTGTTTTGATAGTTGTTCAATT |
| O12 | TCTGTGGCTCCTGAGGCCACACTAGTTAGTGCCAGAAGTGTAAAGACTGCCAAAAACATATAGCGATGAG GCATTGTCATTGTGTTTTGATAGTTGTTCAATT |
| O13 | CAACGCATTCACGCCATTGC |
| O14 | CGAGATATCAGTTTGTCCTCCG |
| O15 | GAAACCGTAACCGATGTGTGG |
| O16 | ACTAACTGCCGGTTTTATTGCAGTTGGCTAAATACGAAAATAATCGGAAATCCACTGCTGCACCCAGTCTT AAATTTTGAAATGTATCTAAACGCAAACTCCGAG |
| O17 | TTGATACCACAAATGGAAACATTTGTATTGACTGTGTCCTCTCAGGCCTTAAAGGAAATTTGGAACTCCAGT GCCACTTTGCACAAACGAAGGTCTCACTTAATC |
| O18 | GCTTCTTTCAAGCAAGCCTTGG |
| O19 | CTATTTGCACACACGAGCGG |
| O20 | CCAGATCATCCCAAGGAACCC |
| O21 | GTTAATTGATCAAAACGTTGGCCC |
| O22 | ATAAACGTATTCAAACTACCCAGTGATGTAGTTATTTATTAAGATTTGAGGGAATGCCACAATGATGCTTCA AGTTACTCAATGTATCTAAACGCAAACTCCGAG |
| O23 | AGATTAACTACAAAACTATACCTACTGTTAGTATCGCATCTTCCAGAATTTTTGAACACCCATAAACACTTGT CATTAAAGCACAAACGAAGGTCTCACTTAATC |
| O24 | CGTATGGTCTCCCACTACACTAC |
| O25 | GTGTTTTTGCAGTTTAGTCGCC |
| O26 | TATAAGGAAGGAAGGCGAAGATCCTGATTTAATCGAATGGACCATTACTGTGTTGTCGCGCACACGACCT AAATACGATCTAACATCCAAAGACGAAAGG |
| O27 | TTAAGCTACACCCTCCTCGTATAATAAACTTGCTCAACTCACAAGCAAACAGTAGTCAACTGAACCCGGTC CTTACTATTCCCACAATTGGGATTAACCAG |
| O28 | ATCAAGGTGGTTAGGTGAATCTTG |
| O29 | GGCCGTTAGCATTTCAACGAACC |
| O30 | GATGACCCGCATGCGATTATG |
| O31 | GAAGTCGAACCTCTAGAGGAAC |
| O32 | GTTGGAATACAGTGCAATCGAAGGGTTGCAGGTTCAATTCCTGTCCGTGTCAAACTTTAATTTTTTCATTTT TTGTTACTAATGTATCTAAACGCAAACTCCGAG |
| O33 | AGAGGACGAGCTTGATACTAAGTTCTAATAGATGTAGCTAAAGTGATTTGATTGATCTTAAAACGTAGAGG TTGGTACTACCCACAATTGGGATTAACCAG |
| O34 | ATCCATAGATACGTGGAACAAATGG |
| O35 | GTTGGAATACAGTGCAATCGAAGGGTTGCAGGTTCAATTCCTGTCCGTGTCAAACTTTAATTTTTTCATTTT TTGTTACTTAATCGAACTCCGAATGCGGTTCTC |
| O36 | GTTGGTATTGTGAAATAGACGCAGATCGGGAACACTGAAAAATAACAGTTATTATTCGAGATCTAACATCC AAAGACGAAAATGTATCTAAACGCAAACTCCG |
| O37 | GGCTCCTGAGGCCACACTAGTTAGTGCCAGAAGTGTAAAGACTGCCAAAAACATATAGCGATGAGGCATT GTCATGATTTGATTGATGAAGGCAGAGAG |
| O38 | CCGTTGATGGAACCGAAAATGTGCAACTCGTGAGTAGCTGGCAAGGAAGCCATGTTGTCCTCTTCACCCT TAGAAACCATGATTTGATTGATGAAGGCAGAGAG |
| O39 | AGCGATATCCTAGTTCTAGGTGG |
| O40 | AACCCACCTAGAACTAGGATATC |

| | |
|---|---|
| O41 | AGCGACAGCAATATATAAACAGA |
| O42 | AACTCTGTTTATATATTGCTGTC |
| O43 | AGCCAACTCGAATTATAGTGGCG |
| O44 | AACCGCCACTATAATTCGAGTTG |
| O45 | AGCTACATGGAATAGGGTCACGT |
| O46 | AACACGTGACCCTATTCCATGTA |
| O47 | AGCCCTAAATACTACCTAAACAG |
| O48 | AACCTGTTTAGGTAGTATTTAGG |
| O49 | AGCGGTTGGTACTATGTCCAACA |
| O50 | AACTGTTGGACATAGTACCAACC |
| O51 | GCGTTACCTTTAATAACCCACACC |
| O52 | GCTCCGCGACGGGGAATTGAAC |
| O53 | ATGCGTTGAAGCGATCCTC |
| O54 | GTTTTAGAGCTAGAAATAGC |
| O55 | TGAAATCAGCGGAGTGGAGG |
| O56 | GTTCAATTCCCCGTCGCGGAGCGGAAGAGCACCCGAAGTATG |
| O57 | TTGCTATTTCTAGCTCTAAAACCGAAGAGCTGCAGACTGGCTG |
| O58 | CGCCCCTTAGATTAGATTGC |
| O59 | GGAGACCAACATGTGAGC |
| O60 | GCCTTTTGCTCACATGTTGGTCTCCaccAAGCTTGCAAATTAAAGC |
| O61 | GCAATCTAATCTAAGGGGCGCTAGAGTGTTGTTACTTTATAC |

### Tab. S2: Plasmids used in this study

| Plasmid name | Description | Reference |
|---|---|---|
| R_C2_gRNAgut1_kan | All-in-one CRISPR plasmid containing Cas9, sgRNA to target the *GUT1* ORF and geneticin/G418 resistance marker | Dalvie *et al.*, 2020 |
| pCas9Gc_ccdB | All-in-one CRISPR plasmid containing Cas9, *ccdB* stuffer with SapI restriction sites for gRNA cloning and geneticin/G418 resistance marker | this study |
| pCas9Hc_ccdB | All-in-one CRISPR plasmid containing Cas9, *ccdB* stuffer with SapI restriction sites for gRNA cloning and Hygromycin resistance marker | this study |
| pCas9Hc_$P_{AOX1}$ | All-in-one CRISPR plasmid containing Cas9, sgRNA to target $P_{AOX1}$ and Hygromycin resistance marker | this study |
| pCas9Hc_*AOX1* | All-in-one CRISPR plasmid containing Cas9, sgRNA to target *AOX1* ORF and Hygromycin resistance marker | this study |
| pCas9Hc_int6 | All-in-one CRISPR plasmid containing Cas9, sgRNA to target the int6 integration site and Hygromycin resistance marker | this study |
| pCas9Hc_int18 | All-in-one CRISPR plasmid containing Cas9, sgRNA to target the int18 integration site and Hygromycin resistance marker | this study |
| pCas9Hc_PNSII-4 | All-in-one CRISPR plasmid containing Cas9, sgRNA to target the PNSII-4 integration site and Hygromycin resistance marker | this study |
| pCas9Hc_PNSI-2 | All-in-one CRISPR plasmid containing Cas9, sgRNA to target the PNSI-2 integration site and Hygromycin resistance marker | this study |
| pBSY5Z | *K. phaffii* expression vector containing the PDF promoter and the transcription terminator of the *K. phaffii AOX1* gene and a Zeocin resistance marker, by cutting with SapI a stuffer between PDF and *AOX1TT* is cut out | bisy GmbH |
| pBSY5Z_NANOBODY® VHH | *K. phaffii* expression vector containing the PDF-aMF-NANOBODY® VHH-*AOX1*TT expression cassette and a Zeocin resistance marker | this study |
| pBB1_mNG_23 | Plasmid containing mNeonGreen (mNG) ORF, serves as template to generate donor DNA | this study |
| pBB1_PGAP_12 | Plasmid containing $P_{GAP}$, serves as template to generate donor DNA | this study |
| pBB1_PDF_12 | Golden Gate basic module containing the PDF promoter | this study |
| pBB1_PDC_12 | Golden Gate basic module containing the PDC promoter (*K. phaffii CTA1*/*CAT1* promoter) | this study |
| pBB1_MIT1_23 | Golden Gate basic module containing the *MIT1* ORF of *K. phaffii* | this study |
| pBB1_MXR1_23 | Golden Gate basic module containing the *MXR1* ORF of *K. phaffii* | this study |
| pBB1_ScRPL3TT_34 | Golden Gate basic module containing the transcription terminator of the *S. cerevisiae RPL3* gene | this study |
| pBB3Hi_095 | Plasmid containing the $P_{AOX1}$-*MIT1*-*ScRPL3*TT expression cassette, serves as template to generate donor DNA | this study |
| pBB3Hi_096 | Plasmid containing the $P_{AOX1}$-*MXR1*-*ScRPL3*TT expression cassette, serves as template to generate donor DNA | this study |
| pBB3Zi_086 | Plasmid containing the PDF-*MIT1*-*ScRPL3*TT expression cassette, serves as template to generate donor DNA | this study |
| pBB3Zi_098 | Plasmid containing the PDC-*MXR1*-*ScRPL3*TT expression cassette, serves as template to generate donor DNA | this study |

### Tab. S3: Yeast strains used in this study

| Strain name | Genotype | Reference/Source |
|---|---|---|
| NRRL Y-11430 | - | United States Department of Agriculture (USDA) and Agricultural Research Service (ARS) |
| NRRL Y-11430 + *HAC1* | NRRL Y-11430::PDC-*HAC1*$_{\text{spliced}}$-*AOX1*TT | De Groeve *et al.*, 2023 |
| yEIA054 | NRRL Y-11430 + *HAC1*<br>int6:PDF-aMF-NANOBODY® VHH-*AOX1*TT | this study |
| yEIA055 | NRRL Y-11430 + *HAC1*<br>int18::PDF-aMF-NANOBODY® VHH-*AOX1*TT | this study |
| yEIA056 | NRRL Y-11430 + *HAC1*<br>int6_int18::PDF-aMF-NANOBODY® VHH-*AOX1*TT | this study |
| yEIA004 | NRRL Y-11430<br>*aox1*Δ::$P_{AOX1}$-*ScFLO1* | this study |
| yEIA010 | NRRL Y-11430 + *HAC1*<br>*aox1*Δ::$P_{AOX1}$-*ScFLO1* | this study |
| yEIA017 | NRRL Y-11430 + *HAC1*<br>*aox1*Δ::$P_{AOX1}$-mNG | this study |
| yEIA018 | NRRL Y-11430 + *HAC1*<br>*aox1*Δ::$P_{GAP}$-*ScFLO1* | this study |
| yEIA019 | NRRL Y-11430 + *HAC1*<br>*aox1*Δ::$P_{GAP}$-mNG | this study |
| yEIA020 | NRRL Y-11430 + *HAC1*<br>*aox1*Δ::$P_{AOX1}$-*ScFLO1*<br>int6::PDF-aMF-NANOBODY® VHH B-*AOX1*TT | this study |
| yEIA024 | NRRL Y-11430 + *HAC1*<br>*aox1*Δ::$P_{AOX1}$-*ScFLO1*<br>int6_int18::PDF-aMF-NANOBODY® VHH B-*AOX1*TT | this study |
| yEIA021 | NRRL Y-11430 + *HAC1*<br>*aox1*Δ::$P_{AOX1}$-mNG<br>int6::PDF-aMF-NANOBODY® VHH B-*AOX1*TT | this study |
| yEIA025 | NRRL Y-11430 + *HAC1*<br>*aox1*Δ::$P_{AOX1}$-mNG<br>int6_int18::PDF-aMF-NANOBODY® VHH B-*AOX1*TT | this study |
| yEIA034 | NRRL Y-11430 + *HAC1*<br>*aox1*Δ::$P_{GAP}$-*ScFLO1*<br>int6::PDF-aMF-NANOBODY® VHH B-*AOX1*TT | this study |
| yEIA038 | NRRL Y-11430 + *HAC1*<br>*aox1*Δ::$P_{GAP}$-*ScFLO1*<br>int6_int18::PDF-aMF-NANOBODY® VHH B-*AOX1*TT | this study |
| yEIA035 | NRRL Y-11430 + *HAC1*<br>*aox1*Δ::$P_{GAP}$-mNG<br>int6::PDF-aMF-NANOBODY® VHH B-*AOX1*TT | this study |
| yEIA039 | NRRL Y-11430 + *HAC1*<br>*aox1*Δ::$P_{GAP}$-mNG<br>int6_int18::PDF-aMF-NANOBODY® VHH B-*AOX1*TT | this study |
| yEIA052 | NRRL Y-11430 + *HAC1*<br>*aox1*Δ::PDF-*ScFLO1*<br>int6_int18::PDF-aMF-NANOBODY® VHH B-*AOX1*TT | this study |
| yEIA053 | NRRL Y-11430 + *HAC1*<br>*aox1*Δ::PDF-mNG<br>int6_int18::PDF-aMF-NANOBODY® VHH B-*AOX1*TT | this study |
| yEIA028 | NRRL Y-11430 + *HAC1*<br>*aox1*Δ::$P_{AOX1}$-*ScFLO1*<br>PNSII-4::$P_{AOX1}$-*MIT1*-*ScRPL3*TT | this study |
| yEIA032 | NRRL Y-11430 + *HAC1*<br>*aox1*Δ::$P_{AOX1}$-mNG<br>PNSII-4::$P_{AOX1}$-*MIT1*-*ScRPL3*TT | this study |
| yEIA029 | NRRL Y-11430 + *HAC1*<br>*aox1*Δ::$P_{AOX1}$-*ScFLO1*<br>PNSII-4::$P_{AOX1}$-*MXR1*-*ScRPL3*TT | this study |
| yEIA033 | NRRL Y-11430 + *HAC1*<br>*aox1*Δ::$P_{AOX1}$-mNG<br>PNSII-4::$P_{AOX1}$-*MXR1*-*ScRPL3*TT | this study |
| yEIA042 | NRRL Y-11430 + *HAC1*<br>*aox1*Δ::$P_{AOX1}$-*ScFLO1*<br>int6_int18::PDF-aMF-NANOBODY® VHH B-*AOX1*TT<br>PNSII-4::$P_{AOX1}$-*MIT1*-*ScRPL3*TT | this study |
| yEIA044 | NRRL Y-11430 + *HAC1*<br>*aox1*Δ::$P_{AOX1}$-mNG<br>int6_int18::PDF-aMF-NANOBODY® VHH B-*AOX1*TT<br>PNSII-4::$P_{AOX1}$-*MIT1*-*ScRPL3*TT | this study |

| | | |
|---|---|---|
| yEIA043 | NRRL Y-11430 + *HAC1*<br>*aox1*Δ::P$_{AOX1}$-*ScFLO1*<br>int6_int18::PDF-aMF-NANOBODY® VHH B-*AOX1*TT<br>PNSII-4::P$_{AOX1}$-*MXR1-ScRPL3*TT | this study |
| yEIA045 | NRRL Y-11430 + *HAC1*<br>*aox1*Δ::P$_{AOX1}$-mNG<br>int6_int18::PDF-aMF-NANOBODY® VHH B-*AOX1*TT<br>PNSII-4::PAOX1-MXR1-ScRPL3TT | this study |
| yEIA047 | NRRL Y-11430 + *HAC1*<br>*aox1*Δ::P$_{AOX1}$-*ScFLO1*<br>int6_int18::PDF-aMF-NANOBODY® VHH B-*AOX1*TT<br>PNSII-4::P$_{AOX1}$-*MXR1-ScRPL3*TT<br>PNSI-2::PDF-*MIT1-ScRPL3*TT | this study |
| yEIA048 | NRRL Y-11430 + *HAC1*<br>*aox1*Δ::P$_{AOX1}$-mNG<br>int6_int18::PDF-aMF-NANOBODY® VHH B-*AOX1*TT<br>PNSII-4::P$_{AOX1}$-*MXR1-ScRPL3*TT<br>PNSI-2::PDF-*MIT1-ScRPL3*TT | this study |
| yEIA049 | NRRL Y-11430 + *HAC1*<br>*aox1*Δ::P$_{AOX1}$-*ScFLO1*<br>int6_int18::PDF-aMF-NANOBODY® VHH B-*AOX1*TT<br>PNSII-4::P*AOX1-MXR1-ScRPL3*TT<br>PNSI-2::PDC-*MXR1-ScRPL3*TT | this study |
| yEIA050 | NRRL Y-11430 + *HAC1*<br>*aox1*Δ::P$_{AOX1}$-mNG<br>int6_int18::PDF-aMF-NANOBODY® VHH B-*AOX1*TT<br>PNSII-4::P*AOX1-MXR1-ScRPL3*TT<br>PNSI-2::PDC-*MXR1-ScRPL3*TT | this study |

**Tab. S4: Template DNA and primers used to generate donor DNA by PCR**

| Donor DNA | Primers | Template | Purpose |
|---|---|---|---|
| *ScFLO1* | O1 & O2 | *S. cerevisiae* genomic DNA | Replacement of *AOX1* by Sc*FLO1* |
| mNG | O7 & O8 | pBB1_PpmNG_23 | Replacement of *AOX1* by mNG |
| $P_{GAP}$ | O10 & O11 | pBB1_PGAP_12 | Replacement of $P_{AOX1}$ by $P_{GAP}$ in strain yEIA017 |
| $P_{GAP}$ | O10 & O12 | pBB1_PGAP_12 | Replacement of $P_{AOX1}$ by $P_{GAP}$ in strain yEIA010 |
| PDF-aMF-NANOBODY® VHH-*AOX1*TT | O16 & O17 | pYPP14F0273 | Integration of expression cassette into int6 locus |
| PDF-aMF-NANOBODY® VHH-*AOX1*TT | O23 & O24 | pYPP14F0273 | Integration of expression cassette into int18 locus |
| $P_{AOX1}$-*MIT1* -*ScRPL3*TT | O26 & O27 | pBB3Hi_095 | Integration of expression cassette into PNSII-4 locus |
| $P_{AOX1}$-*MXR1*-*ScRPL3*TT | O26 & O27 | pBB3Hi_096 | Integration of expression cassette into PNSII-4 locus |
| PDF-*MIT1*-*ScRPL3*TT | O32 & O33 | pBB3Zi_086 | Integration of expression cassette into PNSI-2 locus |
| PDC-*MXR1*-*ScRPL3*TT | O35 & O33 | pBB3Zi_098 | Integration of expression cassette into PNSI-2 locus |
| PDF | O36 & O37 | pBB3Zi_086 | Replacement of $P_{AOX1}$ by PDF in strain yEIA024 |
| PDF | O36 & O38 | pBB3Zi_086 | Replacement of $P_{AOX1}$ by PDF in strain yEIA025 |

**Tab. S5: Combinations of donor DNA and CRISPR-Cas9 plasmids used for transformation of starting strains to generate strains used in this study.** Additionally, PCR primers used for colony PCR or sequencing to verify the introduced modification are indicated. Oligonucleotide sequences are given in Tab. S1.

| Starting strain | Donor DNA | CRISPR-Cas9 plasmid | Resulting strain | PCR primers to verify modification |
|---|---|---|---|---|
| NRRL Y-11430 + *HAC1* | PDF-aMF-NANOBODY® VHH B-*AOX1*TT | pCas9Hc_int6 | yEIA0054 | O18 & O19, O20 & O21 |
| NRRL Y-11430 + *HAC1* | PDF-aMF-NANOBODY® VHH B-*AOX1*TT | pCas9Hc_int18 | yEIA0055 | O25 & O19, O20 & O25 |
| yEIA0054 | PDF-aMF-NANOBODY® VHH B-*AOX1*TT | pCas9Hc_int18 | yEIA0056 | O25 & O19, O20 & O25 |
| NRRL Y-11430 | *ScFLO1* | pCas9Hc_*AOX1* | yEIA004 | O1 & O2 |
| NRRL Y-11430 + *HAC1* | *ScFLO1* | pCas9Hc_*AOX1* | yEIA010 | O1 & O2 |
| NRRL Y-11430 + *HAC1* | mNG | pCas9Hc_*AOX1* | yEIA017 | O6 & O9 |
| yEIA010 | $P_{GAP}$ | pCas9Hc_$P_{AOX1}$ | yEIA018 | O13 & O14 |
| yEIA017 | $P_{GAP}$ | pCas9Hc_$P_{AOX1}$ | yEIA019 | O13 & O15 |
| yEIA010 | PDF-aMF-NANOBODY® VHH B-*AOX1*TT | pCas9Hc_int6 | yEIA020 | O18 & O19, O20 & O21 |
| yEIA020 | PDF-aMF-NANOBODY® VHH B-*AOX1*TT | pCas9Hc_int18 | yEIA024 | O25 & O19, O20 & O25 |
| yEIA017 | PDF-aMF-NANOBODY® VHH B-*AOX1*TT | pCas9Hc_int6 | yEIA021 | O18 & O19, O20 & O21 |
| yEIA021 | PDF-aMF-NANOBODY® VHH B-*AOX1*TT | pCas9Hc_int18 | yEIA025 | O25 & O19, O20 & O25 |
| yEIA018 | PDF-aMF-NANOBODY® VHH B-*AOX1*TT | pCas9Hc_int6 | yEIA034 | O18 & O19, O20 & O21 |
| yEIA034 | PDF-aMF-NANOBODY® VHH B-*AOX1*TT | pCas9Hc_int18 | yEIA038 | O25 & O19, O20 & O25 |
| yEIA019 | PDF-aMF-NANOBODY® VHH B-*AOX1*TT | pCas9Hc_int6 | yEIA035 | O18 & O19, O20 & O21 |
| yEIA035 | PDF-aMF-NANOBODY® VHH B-*AOX1*TT | pCas9Hc_int18 | yEIA039 | O25 & O19, O20 & O25 |
| yEIA024 | PDF | pCas9Hc_$P_{AOX1}$ | yEIA052 | O13 & O14 |
| yEIA025 | PDF | pCas9Hc_$P_{AOX1}$ | yEIA053 | O13 & O15 |
| yEIA010 | $P_{AOX1}$-*MIT1* -*ScRPL3*TT | pCas9Hc_PNSII-4 | yEIA028 | O28 & O29, O30 & O31 |
| yEIA017 | $P_{AOX1}$-*MIT1* -*ScRPL3*TT | pCas9Hc_PNSII-4 | yEIA032 | O28 & O29, O30 & O31 |
| yEIA010 | $P_{AOX1}$-*MXR1* -*ScRPL3*TT | pCas9Hc_PNSII-4 | yEIA029 | O28 & O29, O30 & O31 |
| yEIA017 | $P_{AOX1}$-*MXR1* -*ScRPL3*TT | pCas9Hc_PNSII-4 | yEIA033 | O28 & O29, O30 & O31 |
| yEIA024 | $P_{AOX1}$-*MIT1* -*ScRPL3*TT | pCas9Hc_PNSII-4 | yEIA042 | O28 & O29, O30 & O31 |
| yEIA025 | $P_{AOX1}$-*MIT1* -*ScRPL3*TT | pCas9Hc_PNSII-4 | yEIA044 | O28 & O29, O30 & O31 |
| yEIA024 | $P_{AOX1}$-*MXR1* -*ScRPL3*TT | pCas9Hc_PNSII-4 | yEIA043 | O28 & O29, O30 & O31 |
| yEIA025 | $P_{AOX1}$-*MXR1* -*ScRPL3*TT | pCas9Hc_PNSII-4 | yEIA045 | O28 & O29, O30 & O31 |
| yEIA0043 | PDF-*MIT1*-*ScRPL3*TT | pCas9Hc_PNSI-2 | yEIA047 | O30 & O34, O51 & O34 |

| | | | | |
|---|---|---|---|---|
| yEIA045 | PDF-*MIT1*-*ScRPL3*TT | pCas9Hc_PNSI-2 | yEIA048 | O51 & O34 |
| yEIA0043 | PDC-*MXR1*-*ScRPL3*TT | pCas9Hc_PNSI-2 | yEIA049 | O51 & O34 |
| yEIA045 | PDC-*MXR1*-*ScRPL3*TT | pCas9Hc_PNSI-2 | yEIA050 | O51 & O34 |

**Tab. S6: Quality results for NANOBODY® VHH B produced in Ambr250 bioreactor cultivations.** The quality results of VHH B produced in strain equipped with either the flocculation module (PDF-*ScFLO1* or $P_{GAP}$-*ScFLO1*) or the respective mNG equivalent (PDF-mNG or $P_{GAP}$-mNG) are indicated realtive the VHH B produced in a strain carrying only the modifications required for VHH B production, e. g. carrying no flocculation or mNG module (VHH only = yEIA056). For strains equipped with the flocculation modules (PDF-*ScFLO1* or $P_{GAP}$-*ScFLO1*) and the VHH only reference data from two independent bioreactor cultivations (n = 2) are averaged. For the respective mNG equivalent (PDF-mNG or $P_{GAP}$-mNG) strains, only data from a single fermentation run (n = 1) are respected. Product quality was measured for end of fermentation samples.

| **Product quality attribute [Method]** | **Strain** | | | |
|---|---|---|---|---|
| | **yEIA038** | **yEIA039** | **yEIA052** | **yEIA053** |
| | **$P_{GAP}$-*ScFLO1*** | **$P_{GAP}$-mNG** | **PDF-*ScFLO1*** | **PDF-mNG** |
| Purity [CE-SDS] | 99% | 92% | 94% | 95% |
| Low Molecular Weight species (LMW) [CE-SDS] | 104% | 137% | 129% | 123% |
| Acidic Isoforms [icIEF] | 105% | 113% | 109% | 109% |
| Basic Isoforms [icIEF] | 67% | 67% | 83% | 100% |
| Monomer [HPLC] | 97% | 93% | 91% | 96% |
| High Molecular Weight (HMW) species [HPLC] | 112% | 100% | 111% | 104% |
| Total O-glycosylation [LC-MS] | 109% | 111% | 110% | 109% |
| Missing Disulfide Bonds [RP-HPLC] | 111% | 79% | 93% | 100% |

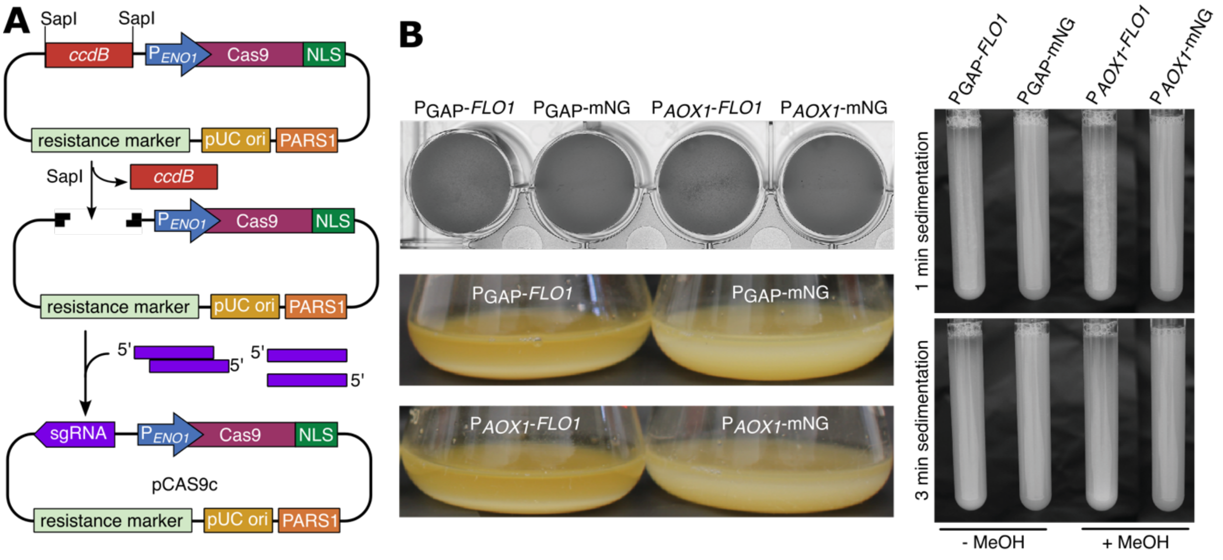


**C**

| Medium | $Ca^{2+}$ conc. (g/L) | $Mg^{2+}$ conc. (g/L) |
|---|---|---|
| 1 | 0.02 | 0.017 |
| 2 | < 0.01 (below test range) | 0.004 |

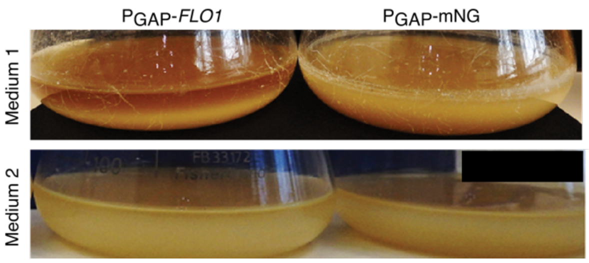


**Fig. S1: *Sc*Flo1p-mediated flocculation assessment**

**A:** Design of CRISPR-Cas9 plasmid. pCas9c vector used for genome editing contains human codon optimized *S. pyogenes* Cas9 fused with SV40 NLS under the control of the $P_{ENO1}$ promoter, a hygromycin resistance cassette, a PARS1 element for auto-replication and a *ccdB* counterselection marker flanked with SapI restriction sites (stuffer) for cloning of sgRNA under the control of a $P_{tRNA1}$–tRNA1 promoter-fusion, which enables transcription and 5' processing of the sgRNA. **B:** Sedimentation behavior of indicated strains after growth in shake flask and MeOH induction. Cells were transferred to culture tubes and photographed after 3 min. For macroscopic analysis, cell suspension was transferred into a well of a 24-well plate and imaged. **C:** Sedimentation behavior of indicated strains after growth in shake flask in different media.

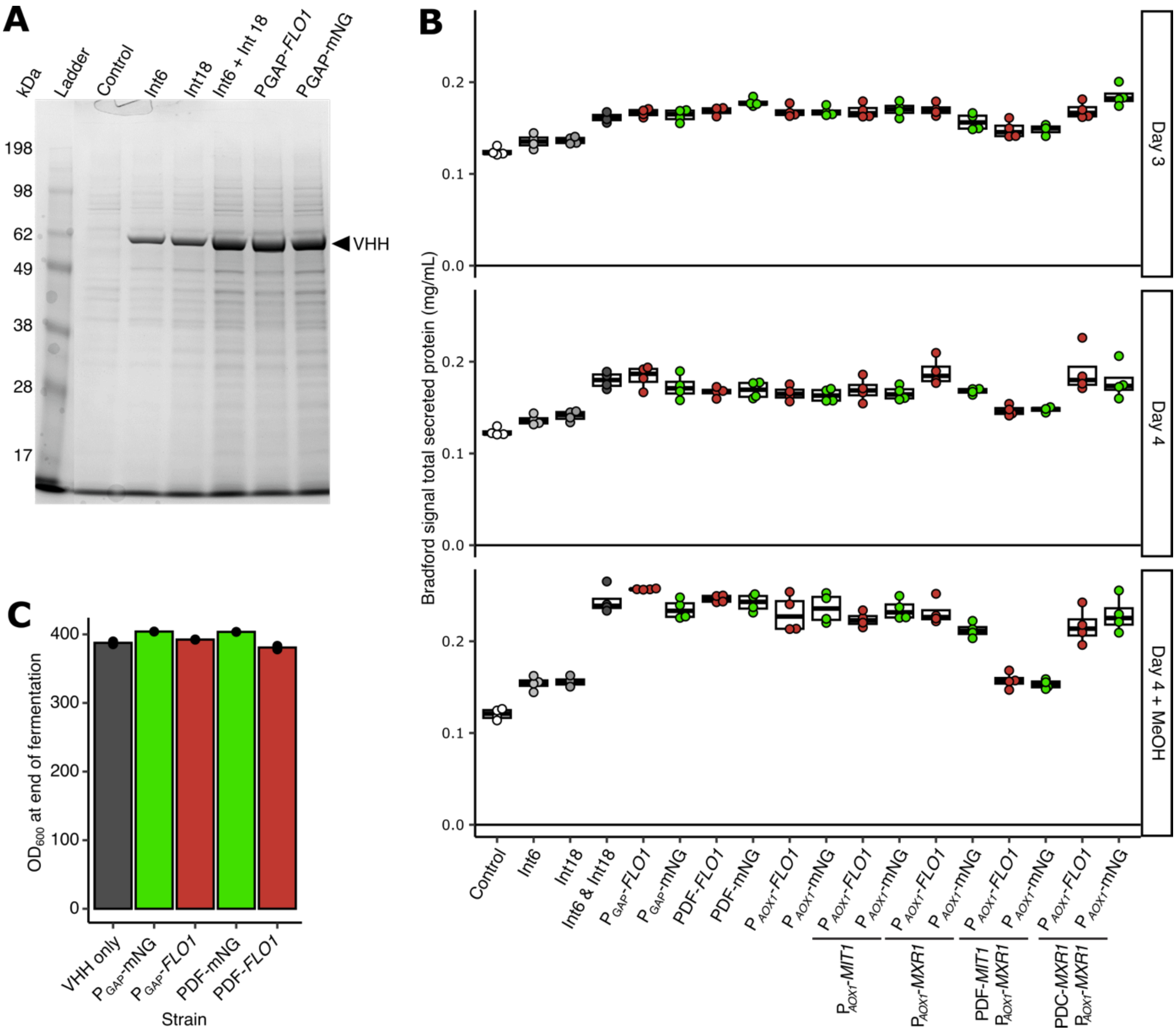


**Fig. S2: Protein secretion in strains with engineered *Sc*Flo1p-mediated flocculation**

**A:** Comparison of NANOBODY® VHH B expression in flocculating strain ($P_{GAP}$-*ScFLO1)* and non-flocculating controls (empty, $P_{GAP}$-mNG*)*. Cell-free supernatant quantification using SDS-PAGE/Coomassie staining. **B:** Total protein quantification in supernatant by semi-automated Bradford assay. Flocculating and non-flocculating control strains were cultivated without and with MeOH addition. **C**: $OD_{600}$ at end of the fermentation runs shown in Fig 4a.

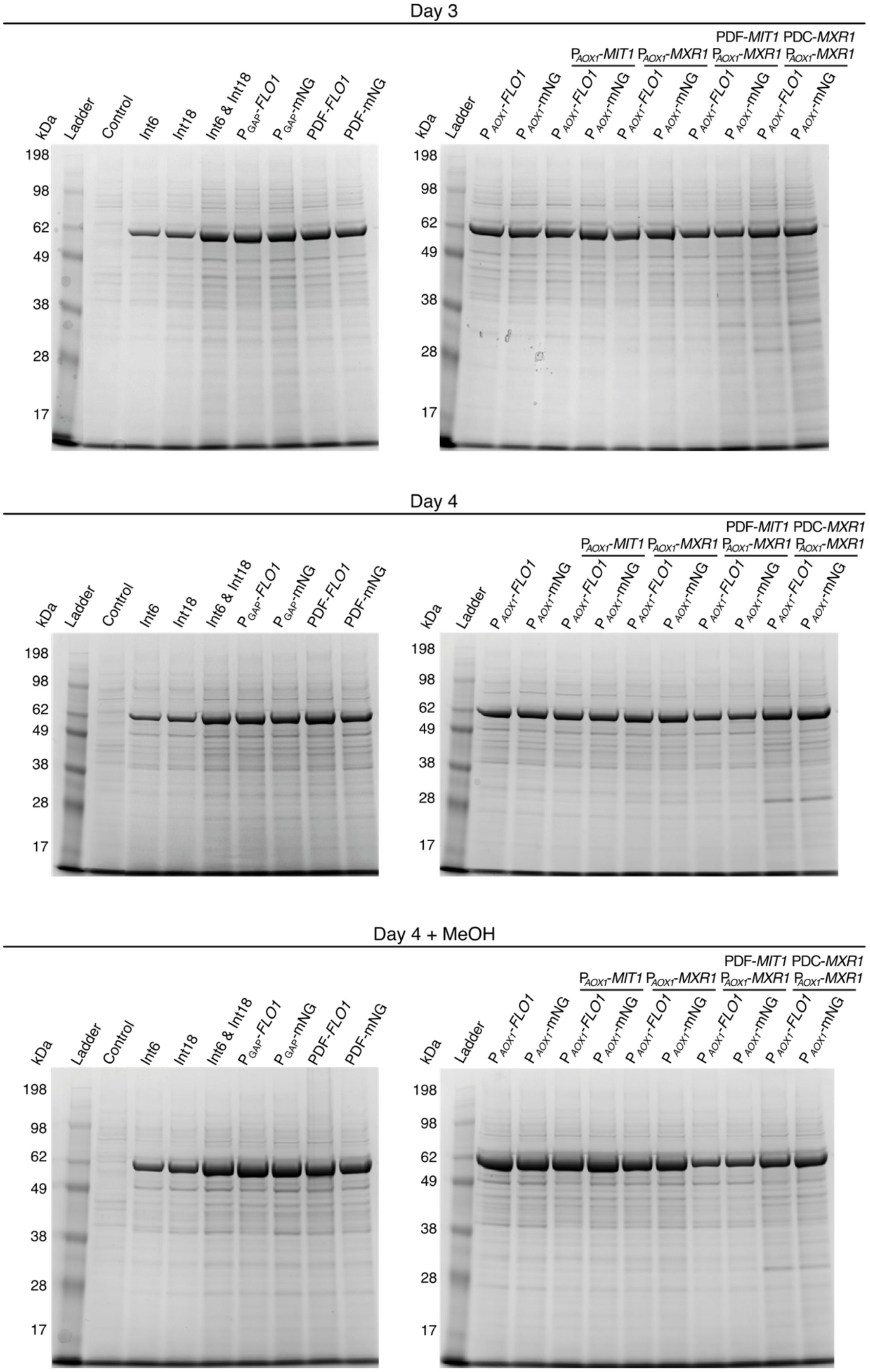


**Fig. S3: Protein secretion in strains with engineered *Sc*Flo1p-mediated flocculation**

Comparison of NANOBODY® VHH B expression in flocculating strains and non-flocculating controls. Flocculating and non-flocculating control strains were cultivated without and with MeOH addition. SDS-PAGE/Coomassie staining of cell-free supernatant.
